# Plastome phylogenomics of the tribe Spermacoceae (Rubiaceae): taxonomic implications and a key to the genera

**DOI:** 10.64898/2026.07.10.737747

**Authors:** Nuñez-Florentin Mariela, Kieran Claypool, Nusrat Huda, Katherine Green, Greg Monzel, Peter W. Schafran, Suman Neupane

**Author notes:** **Corresponding authors:** E-mail addresses (M. Nuñez Florentin), (S. Neupane).

## Abstract

The tribe Spermacoceae (Rubiaceae) comprises a morphologically diverse assemblage of approximately 1,400 species distributed across the Neotropics, Africa, Asia, Australia, and Pacific region. It remains one of the most taxonomically intractable groups in the family, with generic limits repeatedly redefined for more than two centuries. Previous phylogenetic studies based on a limited number of plastid and nuclear markers left numerous relationships unresolved and provided sparse representation of Neotropical lineages. Here, we present the first phylogenomic study of the tribe based on plastome-scale data and expanded sampling of Neotropical taxa. We sampled 121 species representing 55 genera spanning all major clades and generated 123 new plastomes, including 25 species incorporated into a molecular phylogenetic framework for the first time. Maximum-likelihood and Bayesian analyses recovered a highly resolved and strongly supported phylogeny, with uncertainty restricted to a small number of deep backbone nodes. Pollen and seed micromorphology provided additional evidence for evaluating phylogenetic relationships. The resulting phylogenetic framework clarifies generic boundaries across several problematic lineages and supports multiple taxonomic changes. Pervasive homoplasy in seed and floral characters rendered several traditionally recognized genera non-monophyletic, warranting new combinations, including *Edrastima oxycoccoides*, *Stenotis alexanderae*, and *S*. *prostrata*, and a reassessment of taxa such as *Terrellianthus serpyllaceus* and *Oldenlandia dusenii*. We further identify genera requiring additional study and provide an updated key to the 82 recognized genera of Spermacoceae. Together, these results provide the most robust phylogenetic framework yet available for the tribe and establish a foundation for future systematic, biogeographic, and evolutionary research.

## 1. Introduction

Rubiaceae, the fourth-largest family of flowering plants, comprises more than 14,000 species and 570 genera distributed mainly in the tropics and subtropics (Razafimandimbison and Rydin 2024). A growing but still limited number of studies have applied high-throughput genomic data to the family, deepening our understanding of evolutionary relationships at both the subfamily and tribal levels (Ly et al. 2020; Antonelli et al. 2021; Thureborn et al. 2022, 2024; Ball et al. 2023; Thilén et al. 2025). However, several species-rich tribes remain poorly studied from a phylogenomic perspective, particularly those with historically unstable classifications and unresolved generic limits.

Among Rubiaceae tribes, Spermacoceae represents the largest herbaceous lineage, comprising approximately 1,400 species and 84 accepted genera, primarily distributed throughout the tropics and subtropics, particularly in the Neotropics. It is also the most genus-rich and second most species-rich tribe in the family (Verstraete et al. 2025). The tribe has a long and complex taxonomic history, especially at the generic level, because its circumscription has changed repeatedly over the past two centuries (e.g., Berchtold and Presl 1820; Hooker 1873; Schumann 1891; Robbrecht 1988; Andersson and Rova 1999; Wikström et al. 2013; Neupane et al. 2015; Gibbons 2020), and many genera lack clear morphological boundaries for reliable definition. The tribal concept has also varied over time, with up to three different classifications coexisting (Dessein 2003). The currently accepted concept of Spermacoceae (e.g., Kårehed et al. 2008; Groeninckx et al. 2009a), resulted from merging Spermacoceae s.s. with Manettieae and Hedyotideae after molecular studies showed the former nested within the latter (Andersson and Rova 1999). Although previous phylogenetic studies based on a single or a few loci have substantially improved our understanding of Spermacoceae, many deep relationships and generic boundaries remain weakly supported or unresolved, and no phylogenomic study based on high-throughput sequencing data has yet been conducted within the tribe.

Molecular phylogenetic studies on Spermacoceae began with Kårehed et al. (2008) and Groeninckx et al. (2009a), which established the foundational framework for the tribe. These works demonstrated the polyphyly of *Oldenlandia* L. and *Hedyotis* L., confirmed the monophyly of several genera (e.g., *Amphiasma* Bremek.*, Kadua* Cham. & Schltdl., *Phylohydrax* Puff), and revealed complex patterns of para-and polyphyly in others (e.g., *Agathisanthemum* Klotzsch*, Kohautia* Cham. & Schltdl.). Building on this framework, subsequent studies expanded tribal sampling and added new genera through taxonomic discoveries and molecular rearrangements, particularly within African and Asian lineages (Groeninckx et al. 2009b; Groeninckx et al. 2010a, b, c; Guo et al. 2013; Neupane et al. 2015; Xu et al. 2021). More targeted phylogenetic analyses were later conducted to resolve specific taxonomic problems within *Dimetia* (Wight & Arn.) Meisn., *Hedyotis*, *Leptopetalum* Hook. & Arn., among others (Wikström et al. 2013; Neupane et al. 2015; Naiki et al. 2016; Gibbons 2020; Ohi-Toma et al. 2020). Despite these advances, the phylogenetic placement of several taxa, especially among the polyphyletic *Oldenlandia*, remains unclear.

American taxa, particularly South American ones, have been historically underrepresented in these studies. Recent efforts have partially addressed this gap through phylogenetic studies focused mainly on the Spermacoce clade (Salas et al. 2015a; Florentin et al. 2017; Miguel et al. 2018; Nuñez-Florentin et al. 2023, 2024; Carmo et al. 2022, 2024).

Although monophyly was confirmed for several genera (e.g., *Crusea* Cham. & Schltdl., *Galianthe* Griseb., *Richardia* L.), the generic limits of key taxa such as *Borreria* G.Mey., *Spermacoce* L., and *Hexasepalum* Bartl. ex DC., remain unresolved, underscoring the need for broader tribal-level analyses. These limitations have direct consequences for the stability of the taxonomic classification of the tribe, hindering the identification of genera in need of revision or redefinition, and preventing the establishment of a robust evolutionary framework for understanding diversification and biogeographic patterns in the Neotropics.

The present study addresses these shortcomings by providing the first phylogenomic analysis of Spermacoceae based on plastid genomes, incorporating a substantially denser sampling of Neotropical taxa, including genera not previously included in any phylogenetic analyses. Coupled with expanded taxon sampling, the genome-scale information contained in complete or near-complete plastomes provides a robust framework for testing previously unresolved relationships within the tribe. We also compare the plastid-based phylogeny with nuclear-derived trees based on ITS data to evaluate the congruence between cytoplasmic and nuclear signals across the tribe. Along with the phylogeny, we also incorporate morphological seed and pollen data into our study to further clarify the taxonomy and generic limits. Further, we also provide a generic overview and a global key to all recognized genera for the tribe.

## 2. Materials and methods

### 2.1. Taxon sampling and DNA extraction, library preparation, and sequencing

We sampled 121 species representing 53 genera (Fig. 1) across the phylogenetic and geographic breadth of Spermacoceae (Rubiaceae), including representatives of all major clades recognized in previous phylogenetic studies (Supplementary Table 1). *Batopedina pulvinellata* Robbr. was included as an outgroup. The final plastome dataset consisted of 127 accessions representing 121 species. Of these, 123 plastomes were newly generated in this study, whereas four previously published plastomes *(Oldenlandia corymbosa* L. MT767006, *O. brachypoda* DC. MT767007, *O. diffusa* (Willd.) Roxb. MT767008, and *Hedyotis ovata* Thunb. ex Maxim. MK203877) were downloaded from GenBank and incorporated for phylogenetic analyses. Additionally, 25 sampled species lacked previously available molecular data in public databases and are represented here for the first time in a molecular phylogenetic framework.

**Figure 1.**
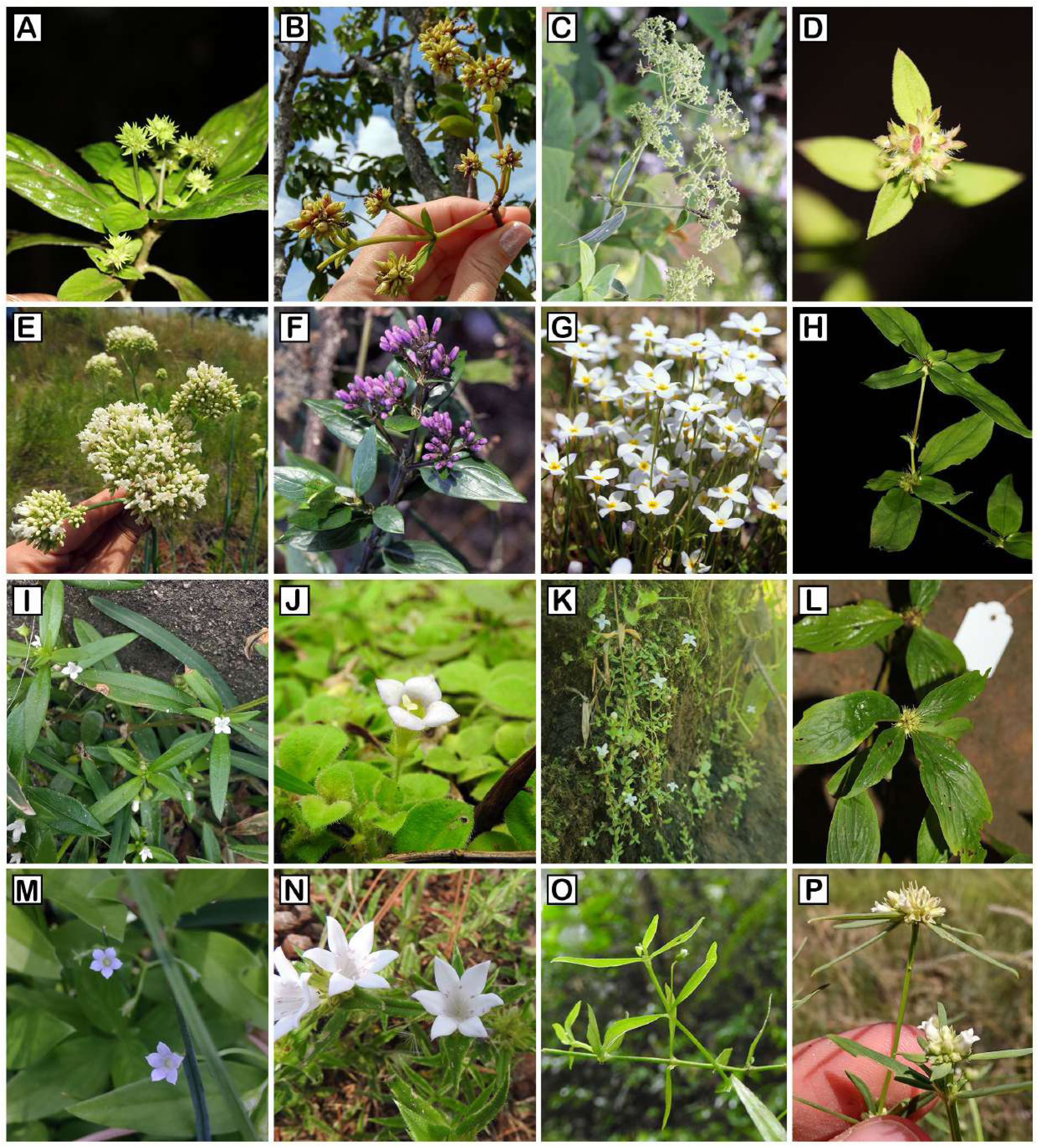
Diversity of Spermacoceae tribe. A. Debia oligocephala. B. Denscantia monodon. C. Dimetia hedyotidea. D. Edrastima uniflora. E. Galianthe fastigiata. F. Hedyotis leschenaultiana. G. Houstonia cearuleae. H. Involucrella coronaria. I. Oldenlandia corymbosa. J. O. dusenii. K. O. salzmannii. L. Parainvolucrella scabra. M. Pentodon pentandrus. N. Richardia grandifolia. O. Scleromitrion lancifolium. P. Spermacoce eryngioides.Photos: A, C–D, F–H, L. Neupane, S. B, E, I–J, M–O. Nuñez-Florentin, M. K, P. Florentin, J.E.

Total genomic DNA was extracted from silica-dried leaf material or herbarium specimens using the DNeasy Plant Mini Kit or DNeasy Plant Pro Kit (Qiagen, Hilden, Germany) following the manufacturer’s protocol. DNA quality and quantity were assessed using agarose gel electrophoresis and a Qubit 4 Fluorometer (Thermo Fisher Scientific).

Sequencing libraries were prepared using either the Nextera XT DNA Library Preparation Kit (Illumina, San Diego, CA, USA) or the NEBNext Ultra II DNA Library Prep Kit for Illumina (New England Biolabs, MA, USA). Libraries were dual-indexed, pooled equimolarly, and sequenced on an Illumina NovaSeq platform using 150 bp paired-end reads, generating genome-skimming data sufficient for plastome assembly.

### 2.2. Plastome assembly, annotation, and sequence alignment

Raw reads were quality-filtered using fastp v0.23.2 (Chen et al. 2018) to trim low-quality bases, remove adapter sequences and poly-G tails associated with Illumina two-color chemistry, and discard reads shorter than 50 bp. Complete plastomes were assembled de novo using NOVOPlasty v4.3.1 (Dierckxsens et al. 2017) and GetOrganelle v1.7.5 (Jin et al. 2020), two assemblers optimized for circular organellar genomes. For samples that failed to produce complete assemblies using the automated pipelines, particularly those with low sequencing coverage or high levels of missing data from herbarium specimens, we employed a reference-guided manual assembly approach. Quality-filtered reads were mapped to a closely related reference plastome using Bowtie2 v2.3.5 (Langmead and Salzberg 2012) with the --very-sensitive-local parameter. Extracted plastid reads were de novo assembled using SPAdes v3.13.0 (Bankevich et al. 2012) with multiple k-mer lengths (21, 33, 55, 77) and the - - careful flag for mismatch and indel correction. Resulting scaffolds and reads were iteratively mapped to the reference in Geneious Prime (https://www.geneious.com) to close assembly gaps, resolve ambiguities in repetitive regions, and verify nucleotide accuracy. Consensus sequences were generated by integrating scaffold-based and read-based assemblies, with manual curation to resolve conflicts and ensure adequate read support across the entire plastome. Gene annotation was performed using GeSeq (Tillich et al. 2017), followed by manual verification of gene boundaries, start and stop codons, and intron positions.

Annotated plastomes were compared across taxa to confirm gene content and order consistency. The three structural partitions [the large single-copy (LSC) region, small single-copy (SSC) region, and one copy of the inverted repeat (IRa)] were extracted and aligned independently using MAFFT v7.490 (Katoh & Standley2013) before concatenation into a single supermatrix.

### 2.3. Whole-plastome phylogenetic analysis

Maximum likelihood (ML) phylogenetic analysis of the whole-plastome dataset was conducted in IQ-TREE 3 (Wong et al. 2025). The concatenated supermatrix comprised the large single-copy (LSC), small single-copy (SSC), and one copy of the inverted repeat (IRa) region to avoid redundancy, representing 127 accessions (121 species) and 134,166 aligned base pairs. Although the supermatrix was structured across three plastome regions, the dataset was analyzed as an unpartitioned matrix with a single best-fit substitution model selected under the Bayesian Information Criterion (BIC) using ModelFinder (Kalyaanamoorthy et al. 2017) with the -m MFP option. Branch support was assessed using ultrafast bootstrap approximation (UFBoot2; Hoang et al. 2018) with 5,000 replicates, and the -bnni option was applied to reduce bootstrap overestimation bias. The same supermatrix was additionally analyzed in a Bayesian framework using PhyloBayes-MPI v1.9 (Lartillot et al. 2013) under the CAT-GTR+Γ model. The CAT component employs a Dirichlet-process mixture (Lartillot and Philippe 2004) to infer site-specific equilibrium-frequency profiles directly from the data, with the number of profile components estimated rather than fixed a priori. These profiles were combined with a single set of general time-reversible (GTR) exchange rates shared across all components, and among-site rate variation was modeled using a discrete gamma distribution with four categories. This site-heterogeneous approach has been shown to reduce systematic biases associated with model misspecification, including long-branch attraction, in large phylogenomic datasets (Lartillot and Philippe 2004; Lartillot et al. 2007). Two independent chains were run in parallel until convergence. Convergence and mixing were assessed using bpcomp, based on the maximum discrepancy across bipartitions (maxdiff < 0.1), and tracecomp, based on effective sample sizes and relative parameter discrepancies.

The first 10% of samples from each chain were discarded as burn-in, and a majority-rule posterior consensus tree was generated from the pooled post-burn-in trees of both chains.

### 2.4. ITS region extraction and phylogenetic analysis

To assess phylogenetic congruence between plastid and nuclear ribosomal datasets, and to evaluate potential cytonuclear discordance in Spermacoceae, the nuclear ribosomal ITS region (ITS1, 5.8S, ITS2) was recovered from the same genome-skimming data used for plastome assembly. nrDNA repeat regions were assembled using GetOrganelle v1.7.5 (Jin et al. 2020) with the embplant_nr database, 10 extension rounds (-R 10), and k-mer sizes of 21, 45, 65, 85, and 105. The longest assembled nrDNA contig per sample was retained for downstream analysis. ITS boundaries were identified and subregions extracted using ITSx v1.1.3 (Bengtsson-Palme et al. 2013), retaining partial detections (--preserve T). The complete ITS region (ITS1, 5.8S, ITS2) was used for subsequent analyses. Extracted sequences were aligned using MAFFT v7.490 (Katoh and Standley 2013). Phylogenetic relationships were inferred using maximum likelihood in IQ-TREE version 3 (Wong et al. 2025) under the GTR+G substitution model.

### 2.5. Micromorphology

In order to test monophyly of specific clades, and morphological convergence, the micromorphology of pollen, fruit, and seeds were analyzed for selected species. For the palynological analyses, pollen grains were acetolysed according to the technique by Erdtman (1966) and mounted on glycerine jelly for light microscopy (LM) analysis, under a Leica DM LB2 OM equipped with a digital camera. Additionally, pollen grains as well as fruit and seeds micro-morphology were photographed with a scanning electron microscope (SEM, Zeiss Evo15 from CME-UNNE: Centro de Microscopía Electrónica de la Universidad Nacional del Nordeste). For the SEM observation, pollen grains were mounted on gold-plated aluminium plates, and fruits and seeds were mounted on aluminium stubs with double-sided adhesive tape without any treatment. In both cases, the materials were sputter-coated with 20 nm of gold-palladium. The scaled photos were measured using ImageJ software (Rasband 2020). At least 20 pollen grains from each exsiccate analyzed were measured. Information on species and specimens included in the micromorphological analyses is available in Supplementary Table 2. Pollen terminology followed Punt et al. (2007), and seeds and fruit terminology followed Stearn (1986), Terrell and Robinson (2007).

## 3. Results

### 3.1. Chloroplast genome features

Among the 123 newly generated plastomes, 73 were recovered as complete circular assemblies directly from NOVOPlasty and/or GetOrganelle, requiring no manual scaffolding. The remaining 50 plastomes were assembled through manual scaffolding of partial assemblies and are described separately (see section 2.2). The 73 fully assembled plastomes characterized here all displayed the canonical quadripartite structure, comprising a large single-copy region (LSC, 81,776–85,249 bp), a small single-copy region (SSC, 16,831–19,161 bp), and two inverted repeat regions (IRs, 24,963–26,376 bp). Total plastome length ranged from 151,100 bp in *Terrellianthus serpyllaceus* (Schltdl.) Borhidi to 155,066 bp in *Kohautia cynanchica* DC., with a mean of 153,114 bp. Overall GC content was highly conserved, averaging 37.6% (range: 37.1–38.0%). Gene content was broadly conserved across sampled taxa, with most plastomes containing approximately 79–80 unique protein-coding genes and ∼30 tRNA genes, together with the typical plastid rRNA complement, for a total of 112–118 annotated unique genes. No evidence of large-scale gene loss or major structural rearrangement was detected. Most variation was limited to modest differences in LSC, SSC, and IR lengths, consistent with minor IR boundary expansion and contraction.

Annotated lengths of several short or rapidly evolving genes (e.g., infA, ycf15) varied across taxa, although detailed characterization of plastid gene evolution in Spermacoceae is beyond the scope of the present study.

### 3.2. Plastome molecular phylogeny

The final plastome matrix comprised 127 plastome sequences representing 121 species and 56 genera (including *Batopedina* as the outgroup), with 134,166 aligned nucleotide sites from the large single-copy (LSC), small single-copy (SSC), and one copy of the inverted repeat (IRa) region. The LSC region contributed the largest proportion of the alignment (88,687 sites), followed by IRa (26,400 sites) and SSC (19,079 sites). Across the complete matrix, 34,462 sites were parsimony-informative, representing 25.7% of the alignment, whereas 80,836 sites were constant. The majority of parsimony-informative sites were located in the LSC region (25,713 sites), followed by SSC (6,796 sites), while IRa was comparatively conserved, containing only 1,953 parsimony-informative sites. Maximum likelihood (ML, Figure 2A–B) and Bayesian analyses (Supplementary figure 1) recovered a highly resolved and largely congruent plastome phylogeny, with most nodes receiving strong support in both analyses (BS ≥ 95, BPP ≥ 0.95). Bayesian inference under the CAT-GTR+Γ model in PhyloBayes yielded a topology and support pattern that were nearly identical to those recovered in the ML analysis (Supplementary figure 2). Further results will be given primarily based on ML phylogeny (Figure 2A–B).

**Figure 2.**
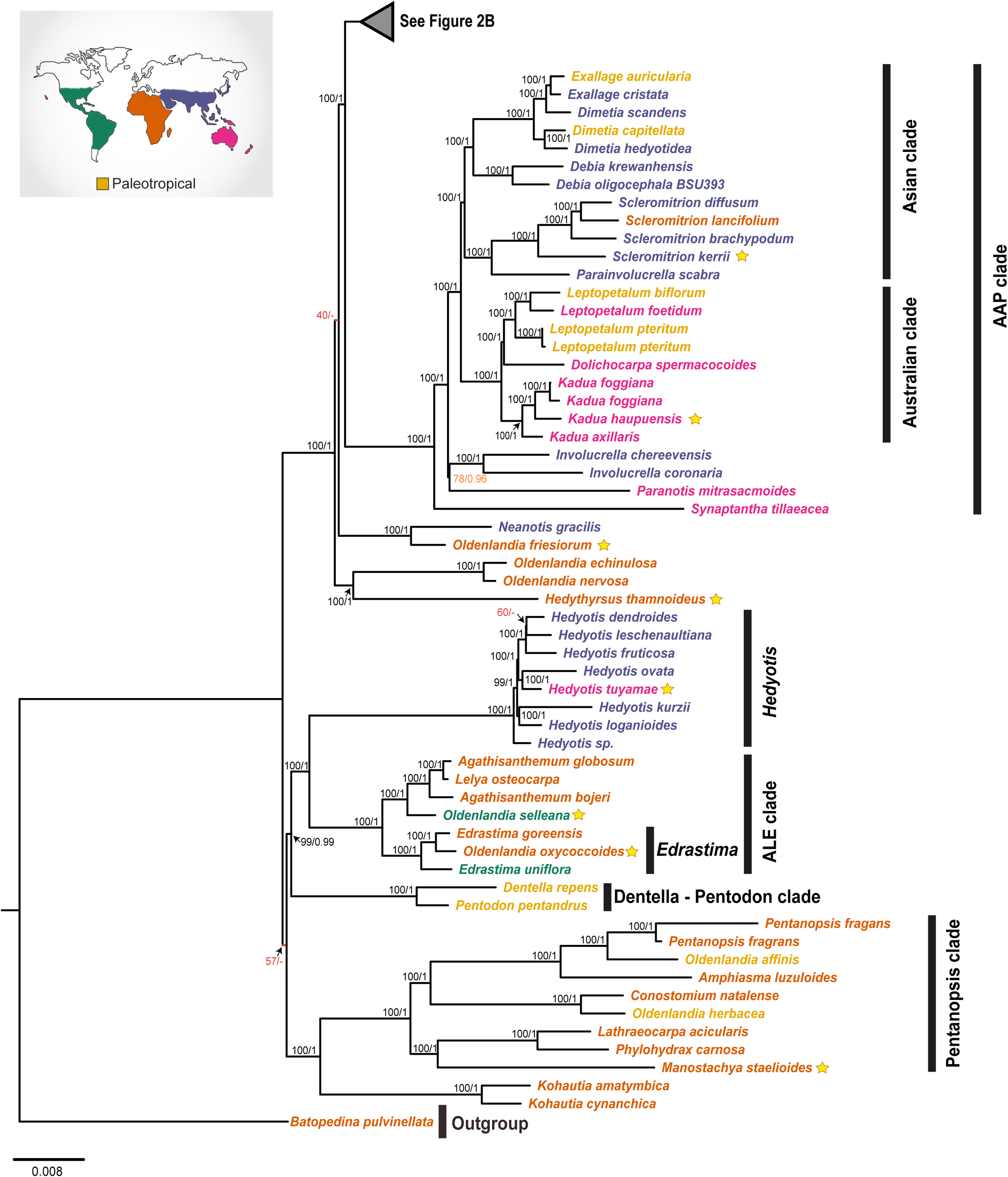
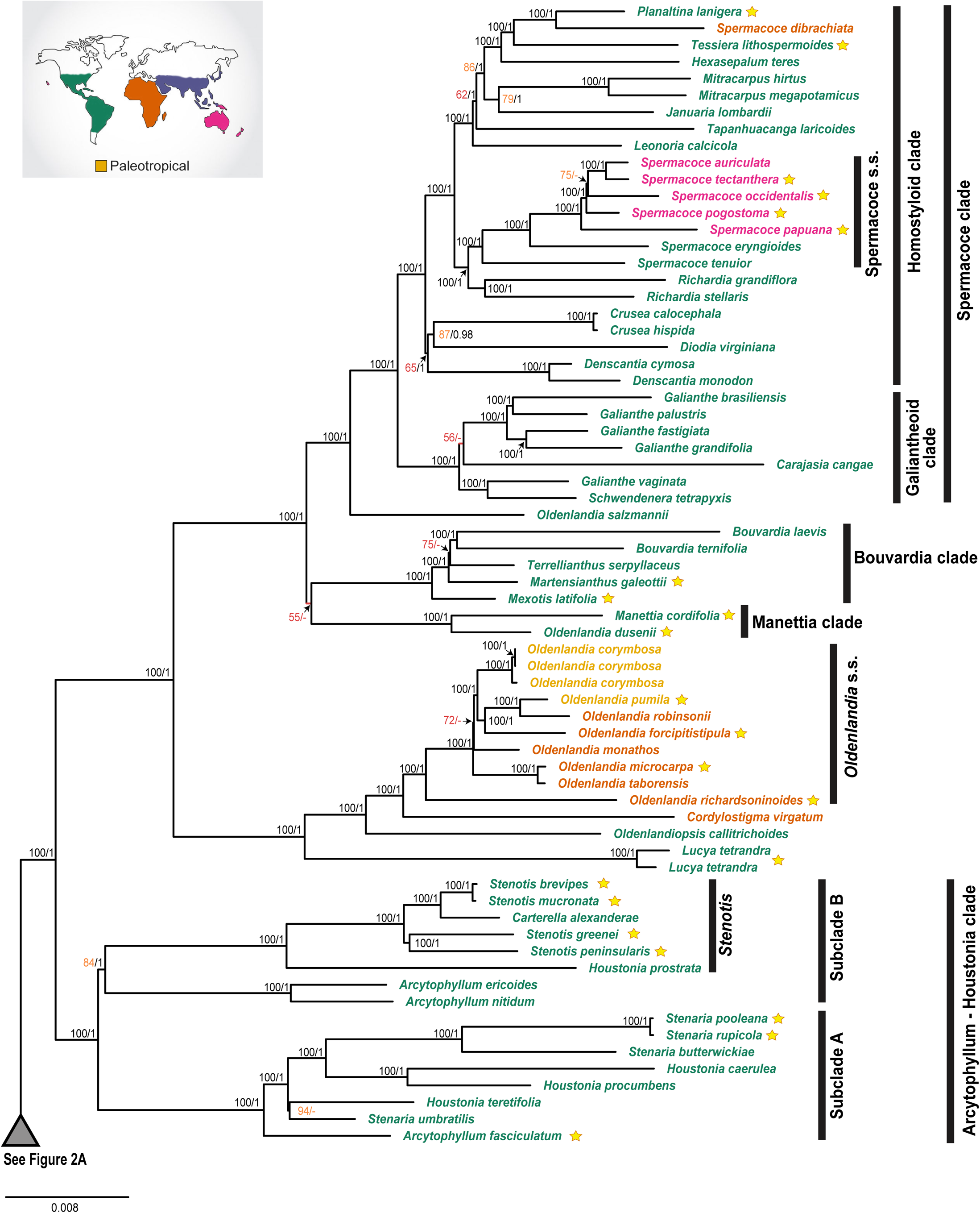
Phylogenetic relationships within tribe Spermacoceae inferred from maximum likelihood (ML) and Bayesian inference (BI) analyses of plastome data. The tree is presented in two parts (A and B). Numbers on the node indicate ML bootstrap support (left) and Bayesian Posterior Probabilities (right). Nodes with BS values = 100 and 1 PP are in black (strong support). Nodes with BS values below 75 are in red (weak support). Nodes with BS values among 75–94 are in orange (moderate support). Values below 0.95 PP are omitted and represented by a dash (-). Species included for the first time in a phylogenetic analysis are indicated with a yellow star. Terminal taxa are colored according to their native biogeographic origin: America (green), Africa (orange), Asia (purple), Australia + Pacific Islands (pink). Taxa distributed across two or more Old World regions (Africa, Asia, and/or Australia + Pacific Islands) are classified as Paleotropical (yellow) and shown in yellow.

The major clades resolved (BS = 100, BPP = 1) are Dentella-Pentodon, Agathisanthemum/Lelya-Edrastima (ALE) clade, *Hedyotis*, Kohautia-Pentanopsis, Asian-Australian-Pacific (AAP) clade, Arcytophyllum-Stenotis-Houstonia/Stenaria), *Oldenlandia* s.s., Bouvardia clade, and Spermacoce clade. Although the primary lineages recovered here are consistent with previous phylogenetic studies, expanded taxon sampling and plastome-scale data provide greater resolution of interclade relationships and the placement of previously unsampled taxa.

*Dentella repens* (L.) J.R.Forst. & G.Forst. and *Pentodon pentandrus* (Schumach.) Vatke were recovered as a strongly supported clade (Fig. 2A), sister to Hedyotis-ALE clade (BS = 99, BPP = 0.99), despite being separated by an extremely short internal branch. The ALE clade (*Agathisanthemum/ Lelya*-*Edrastima* clade) is recovered as a sister to *Hedyotis* with maximum support. *Hedyotis* is recovered as a strongly supported monophyletic group. Internal relationships among the sampled species were subtended by short branches, indicating limited plastome divergence among closely related lineages. The ALE clade itself is highly supported. Within it, *Agathisanthemum* is not recovered as monophyletic, as *Lelya osteocarpa* Bremek. is nested within it. The Hispaniolan endemic *Oldenlandia selleana* Urb., previously unsampled in phylogenetic studies, is recovered as sister to the Agathisanthemum-Lelya clade. Likewise, the pantropical *Edrastima* Raf. is not recovered as monophyletic; the East African *Oldenlandia oxycoccoides* Bremek., also previously unsampled, is instead recovered nested within *Edrastima*. The principal remaining areas of uncertainty were restricted to a small number of deep backbone relationships within core Spermacoceae, including the placement of the Dentella-Pentodon clade *+ Hedyotis +* ALE clade relative to the Pentanopsis clade + *Kohautia* assemblage and the remainder of the tribe.

*Kohautia* is recovered as monophyletic and sister to the Pentanopsis clade. Within this clade, *Manostachya staelioides* (K.Schum.) Bremek. resolved sister to a highly supported clade comprising *Phylohydrax carnosa* (Hochst.) Puff and *Lathraeocarpa acicularis* Bremek.

*Oldenlandia echinulosa* K.Schum. and *O*. *nervosa* Hiern formed a maximally supported clade, with *Hedythyrsus thamnoideus* (K.Schum.) Bremek. resolved as their sister taxon. In contrast, the newly sampled, African *O. friesiorum* Bremek. is strongly supported as sister to the Asiatic *Neanotis gracilis* (Hook.f.) W.H.Lewis. Relationships among these clades and the remainder of the core tribe remained unresolved.

The AAP clade refers to a group of 25 taxa distributed mainly across the Asian-Australo-Pacific region. *Synaptantha tillaeacea* (F.Muell.) Hook.f., endemic to Australia, is recovered as sister to the remaining taxa of this clade. The sister relationship between *Paranotis mitrasacmoides* (F.Muell.) K.L.Gibbons and the Involucrella clade was subtended by an extremely short branch and was weakly supported under maximum likelihood (BS = 78), though moderately supported under Bayesian inference (BPP = 0.96). This assemblage is in turn sister to a highly supported clade comprising *Debia* Neupane & N.Wikstr., *Dimetia*, *Dolichocarpa* K.L.Gibbons, *Exallage* Bremek., *Kadua*, *Leptopetalum*, and *Scleromitrion* (Wight & Arn.) Meisn. Within this larger clade, two strongly supported subclades are identified: the Asian clade and the Australian-Pacific clade. Within the Asian clade, *Scleromitrion* is recovered as monophyletic and sister to a clade formed by *Debia, Dimetia*, and *Exallage*. *Debia* and *Exallage* are each recovered as monophyletic, whereas *Dimetia* is not; *Dimetia scandens* (Roxb.) R.J.Wang is recovered as sister to *Exallage* species. Within the Australia-Pacific clade, *Kadua* is recovered as monophyletic and sister to *Leptopetalum* + *Dolichocarpa spermacocoides* (F.Muell.) K.L. Gibbons.

This AAP clade is recovered as sister to the remaining recognized clades, most of which have an American distribution (Fig. 2B), except the *Oldenlandia* s.s. clade, which is composed mainly of African taxa. The Arcytophyllum-Houstonia clade is recovered with maximum support and is sister to all remaining clades. Two main subclades are recognised with high support: subclade A and subclade B, within which *Arcytophyllum* Schult. & Schult.f., *Houstonia* Gronov., *Stenaria* (Raf.) Terrell, and *Stenotis* Terrell are not recovered as monophyletic. Within subclade A, *Arcytophyllum fasciculatum* (A.Gray) Terrell & H.Rob. is recovered as a sister to a clade composed of *Houstonia* and *Stenaria* species. *Stenaria umbratilis* (B.L.Rob.) Terrell is closely related to *H*. *teretifolia* Terrell (BS = 94), though the branch subtending this relationship is very short. The remaining *Stenaria* species are all recovered together in a strongly supported clade. Within subclade B, *A. ericoides* (Willd.) Standl + *A. nitidum* (Kunth) Schltdl. are recovered as sister taxa to a clade formed by *H. prostrata Brandegee*, *Carterella* Terrell, and *Stenotis* species. *Carterella alexanderae* (A.M.Carter) Terrell appears nested within *Stenotis*, making the latter paraphyletic.

*Lucya tetrandra* (L.) K.Schum., endemic to the Caribbean Islands, is recovered as sister to the Oldenlandia s.s. + *Cordylostigma* + *Oldenlandiopsis* assemblage.

*Oldenlandiopsis callitrichoides* (Griseb.) Terrell & W.H.Lewis another Neotropical taxon results sister to *Cordylostigma* Groeninckx & Dessein, which in turn turns out to be sister to *Oldenlandia* s.s. clade. The whole *Oldenlandia* s.s. + *Cordylostigma* + *Oldenlandiopsis + Lucya* clade is recovered with high support as sister to a clade encompassing the Manettia clade, Bouvardia clade, and Spermacoce clade.

The placement of the *Manettia* clade relative to the *Bouvardia* clade and the remaining sampled Spermacoceae clade (primarily New World Spermacoceae lineages) differed between the ML and Bayesian analyses (Fig. 2B; supplementary Figure 1). Both analyses strongly supported a sister relationship between *Oldenlandia dusenii* Standl. and *M*. *cordifolia* Mart. However, in the ML phylogeny (Fig. 2B), this clade was recovered as sister to the Bouvardia clade (comprising *Bouvardia* Salisb., *Martensianthus* Borhidi & Lozada-Pérez, *Mexotis* Terrell & H.Rob., and *Terrellianthus* Borhidi), although support for this relationship is weak (BS = 55). In contrast, the Bayesian consensus tree (Supplementary Figure 2) placed the *Manettia* Mutis ex L. clade as sister to the Spermacoce clade + *O*. *salzmannii* assemblage (BPP = 0.97), with the Bouvardia clade recovered as the sister to the rest. However, the branch subtending the Manettia clade in the Bayesian topology (Supplementary figure 1) is extremely short, suggesting limited phylogenetic signal for this deep split.

Within the Bouvardia clade (Fig. 2B), *Mexotis latifolia* (M.Martens & Galeotti) Terrell & H.Rob. was resolved as sister to all remaining members, with *Martensianthus galeottii* (M.Martens) Borhidi & Lozada-Pérez in turn sister to the rest of the clade.

*Terrellianthus serpyllaceus* is recovered as sister to *Bouvardia* (clade comprising *B*. *laevis* M.Martens & Galeotti and *B*. *ternifolia* (Cav.) Schltdl.), although support for the *Terrellianthus* + *Bouvardia* is moderate (BS = 75).

*Oldenlandia salzmannii* (DC.) Benth. & Hook.f. ex B.D.Jacks. is recovered as a sister to the Spermacoce clade. Within this clade, the Galiantheoid clade and the Homostyloid clade are each recovered with maximum support. The former includes *Carajasia* R.M.Salas, E.L.Cabral & Dessein, *Schwendenera* K.Schum., and *Galianthe*, with the latter recovered as polyphyletic. *Schwendenera tetrapyxis* K.Schum. + *Galianthe vaginata* E.L.Cabral & Bacigalupo are recovered as highly supported taxa, sister to an assemblage composed of *Carajasia cangae* R.M.Salas, E.L.Cabral & Dessein + remaining *Galianthe* species, although with a weak support (BS = 56). The Homostyloid clade encompasses all the remaining genera. *Crusea, Denscantia* E.L.Cabral & Bacigalupo*, Mitracarpus* Zucc., and *Richardia* are recovered as monophyletic, while *Spermacoce* is polyphyletic. The remaining genera are each represented by a single species, except *Leonoria* Nuñez Florentin & R.M.Salas and *Januaria* R.M.Salas & Nuñez Florentin, which are monospecific. *Denscantia* + *Diodia virginiana* L. + *Crusea* form a clade with moderate support (BS = 65), although it has strong support (BPP = 1) in the Bayesian analysis (Supplementary Figure 1). Within Spermacoce s.s, all Australian and Pacific Islands species are recovered together in a strongly supported clade. In contrast, the African *S. dibrachiata* Oliv. is not recovered among the remaining *Spermacoce* species, appearing instead as sister to *Planaltina lanigera* (DC.) R.M.Salas & E.L.Cabral.

### 3.3. Nuclear vs plastome phylogeny

Figure 3 presents a comparison between nuclear ITS (ITS1, 5.8S, ITS2) and plastome phylogenies. The ITS phylogeny recovered a topology largely congruent with that obtained from the whole-plastome dataset, with most major clades identified in the plastome tree also recovered in the nuclear tree. However, support values were generally lower for both the deepest and shallowest nodes in the ITS phylogeny. The position of *Dentella repens* + *Pentodon pentandrus*, the monophyly of *Hedyotis*, the ALE clade, the Arcytophyllum-Houstonia clade, and the AAP clade are consistently recovered across both trees, supporting their robustness. Similarly, the Spermacoce clade and its two main subclades (Galiantheoid and Homostyloid) are recovered in both trees.

**Figure 3.**
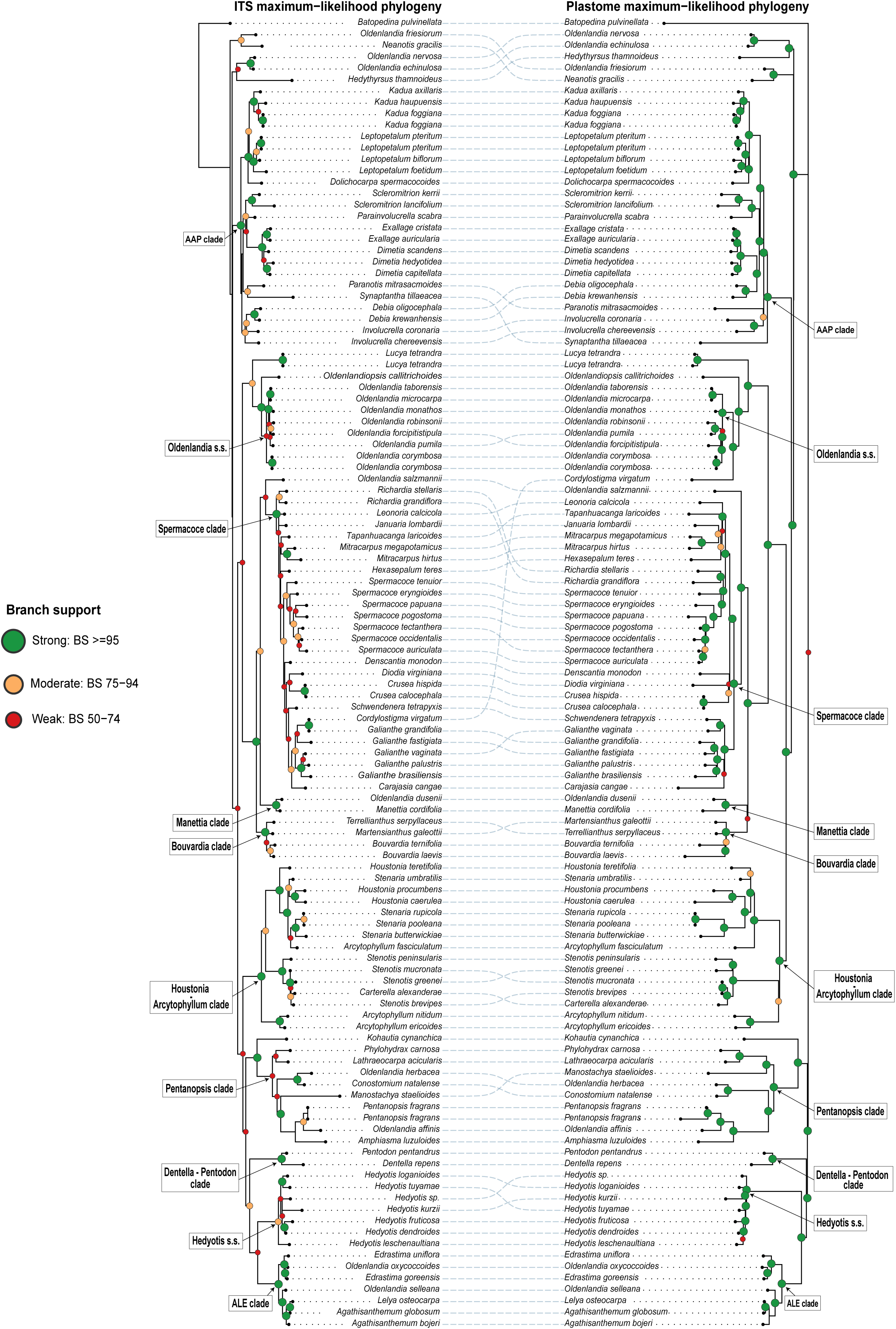
Tanglegram comparing the ITS (left) and plastome (right) maximum-likelihood phylogenies of Spermacoceae. Branch support values are indicated by filled circles at nodes: green circles, BS≥ 95 (strong support); orange circles, BS 75–94 (moderate support); red circles, BS 50–74 (weak support); nodes with BS < 50 are unmarked. Dotted lines connect corresponding taxa across both topologies.

However, several notable inconsistencies are observed between the two topologies. The most evident differences concern the relative arrangement of some mid-level clades, particularly within the AAP clade, where the positions of *Paranotis mitrasacmoides* and *Synaptantha tillaeacea* differ between trees. Within the Arcytophyllum-Houstonia clade, the ITS tree shows reduced support for several relationships that are strongly supported in the plastome phylogeny. The Spermacoce clade also exhibits several topological discordances between trees, with the ITS tree placing some taxa in alternate positions with weak to moderate support (eg., *Richardia* spp., *Tapanhuacanga laricoides*). Strikingly, *Cordylostigma virgatum* is recovered with high support in different positions in the ITS and plastome phylogenies. In general, nodes with high BS values (≥95) are more frequent and more broadly distributed across the plastome phylogeny, whereas moderate (75–94) and low (50–74) support values are proportionally more common in the ITS tree, particularly at internal nodes subtending major clades.

### 3.4. Micromorphology of pollen and seeds

In order to test the monophyly of specific clades, morphological convergence and support phylogenetic relationships, pollen grains of *Carterella alexanderae*, *Lucya tetrandra*, *Martensianthus galeottii*, *Mexotis latifolia*, *Oldenlandiopsis callitrichoides*, and *Terrellianthus serpyllaceus* were examined (Figs. 4–5). All examined taxa present isopolar and zonoaperturate pollen grains, with three colpi in most species, six in *L*. *tetrandra*, and (7-)8(−9) in *O*. *callitrichoides*. The endoaperture is lalongate in all taxa except *L*. *tetrandra* (Fig. 4J) and *O*. *callitrichoides* (Fig. 5O), which bear an endocingulum. All heterostylous taxa exhibit consistent pollen dimorphism, with short-styled flowers producing larger grains than long-styled flowers (Table 1). Further details about the pollen morphology are summarized in Table 1.

**Figure 4.**
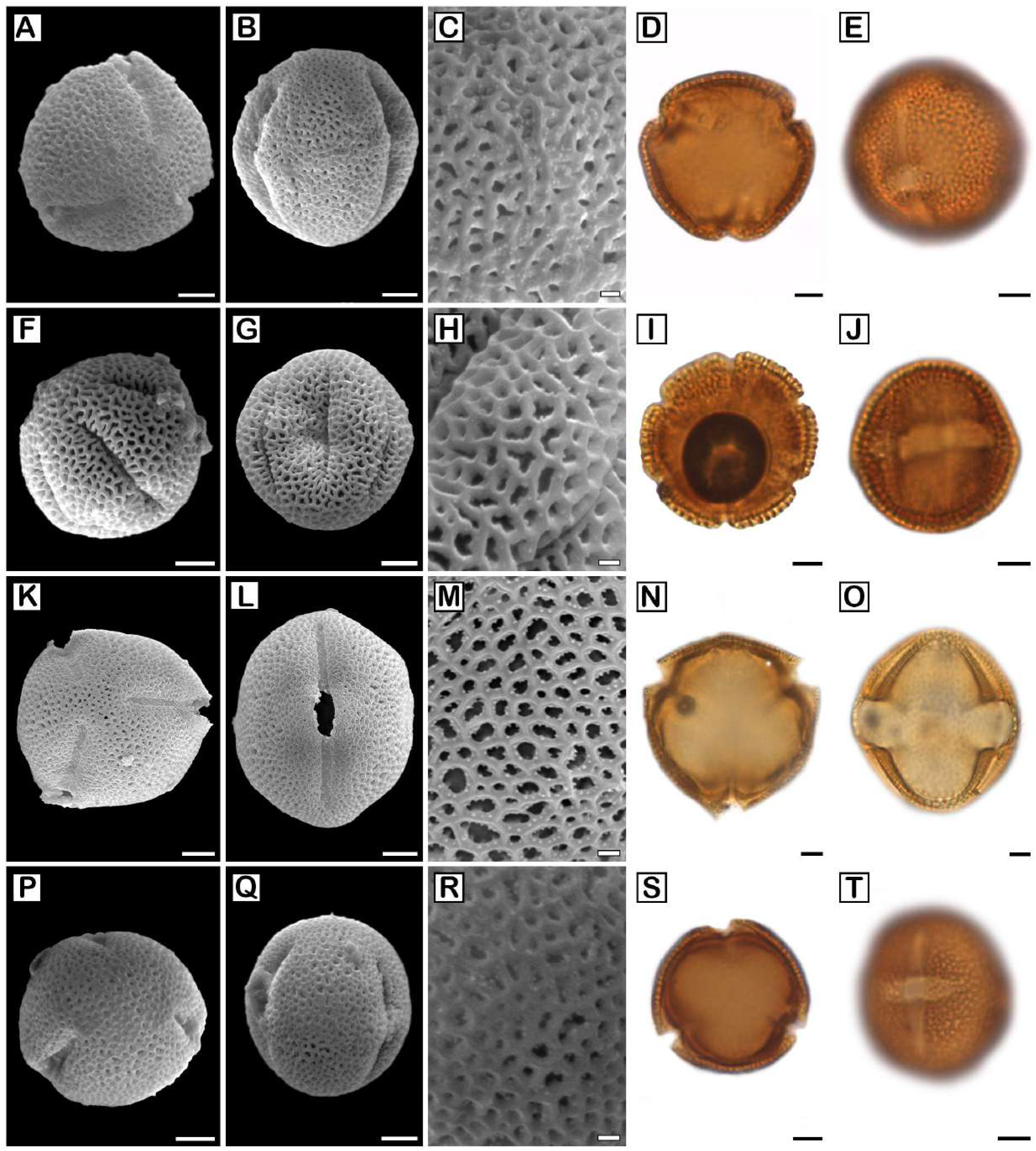
Pollen morphology of selected Spermacoceae taxa. A–C, F–H, K–M, P–R. SEM photographs. D–E, I–J, N–O, S–T. LM photographs. A–E. *Carterella alexanderae* (short-styled flower). F–J. *Lucya tetandra*. K–T. *Martensianthus galeottii*. K–O. Short-styled flower. P–T. Long-styled flower. A, F. Subpolar view. B, G, J, L, O, Q, T. Equatorial view. D, I, K, N, P, S. Polar view. C, H, M, R. Detail of exine. Scale bars: A–B, D–E, F–G, I–J, K–L, N–O, P–Q, S–T. 5 µm. C, H, M, R. 1 µm.

**Figure 5.**
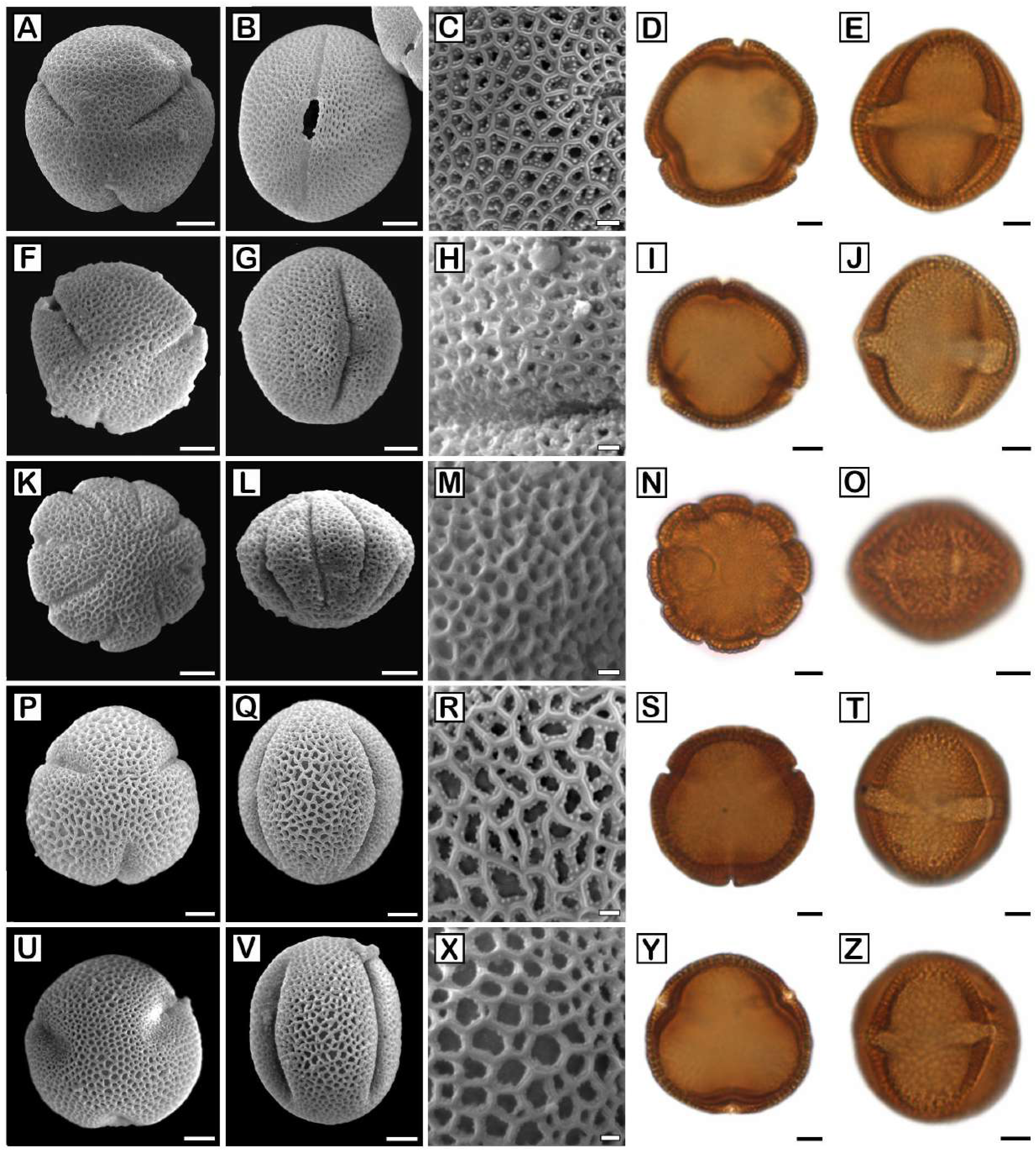
Pollen morphology of selected Spermacoceae taxa. A–C, F–H, K–M, P–R, U–X. SEM photographs. D–E, I–J, N–O, S–T, Y–Z. LM photographs. A–J. *Mexotis latifolia*. A–E. Short-styled flower. F–J. Long-styled flower. K–O. *Oldenlandiopsis callitrichoides*. P–Z. *Terrellianthus serpyllaceus*. P–T. Short-styled flower. U–Z. Long-styled flower. A, D, F, I, K, N, P, S, U, Y. Polar view. B, E, G, J, L, O, Q, T, V, Z. Equatorial view. C, H, M, R, X. Detail of exine. Scale bars: A–B, D–E, F–G, I–J, K–L, N–O, P–Q, S–T, U–V, Y–Z. 5 µm. C, H, M, R, X. 1 µm.

**Table 1.**
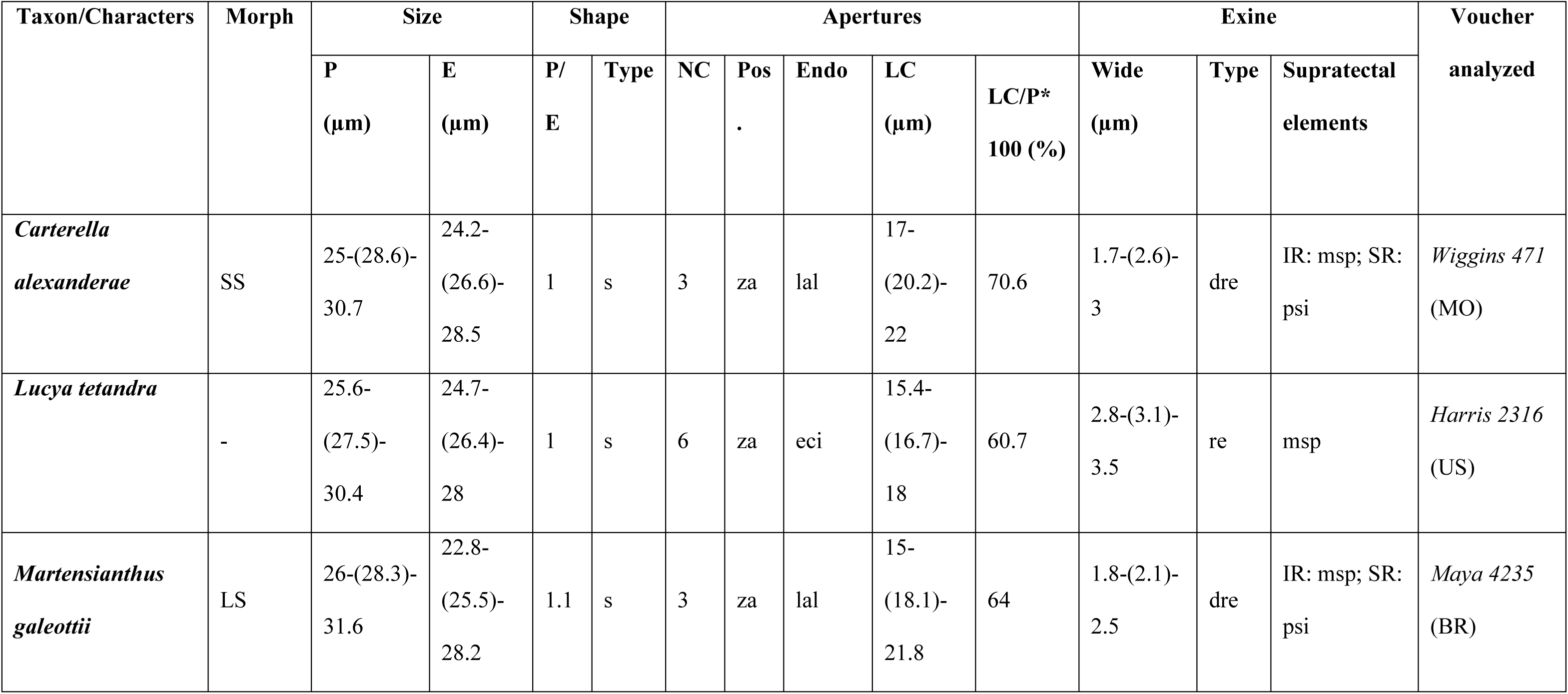
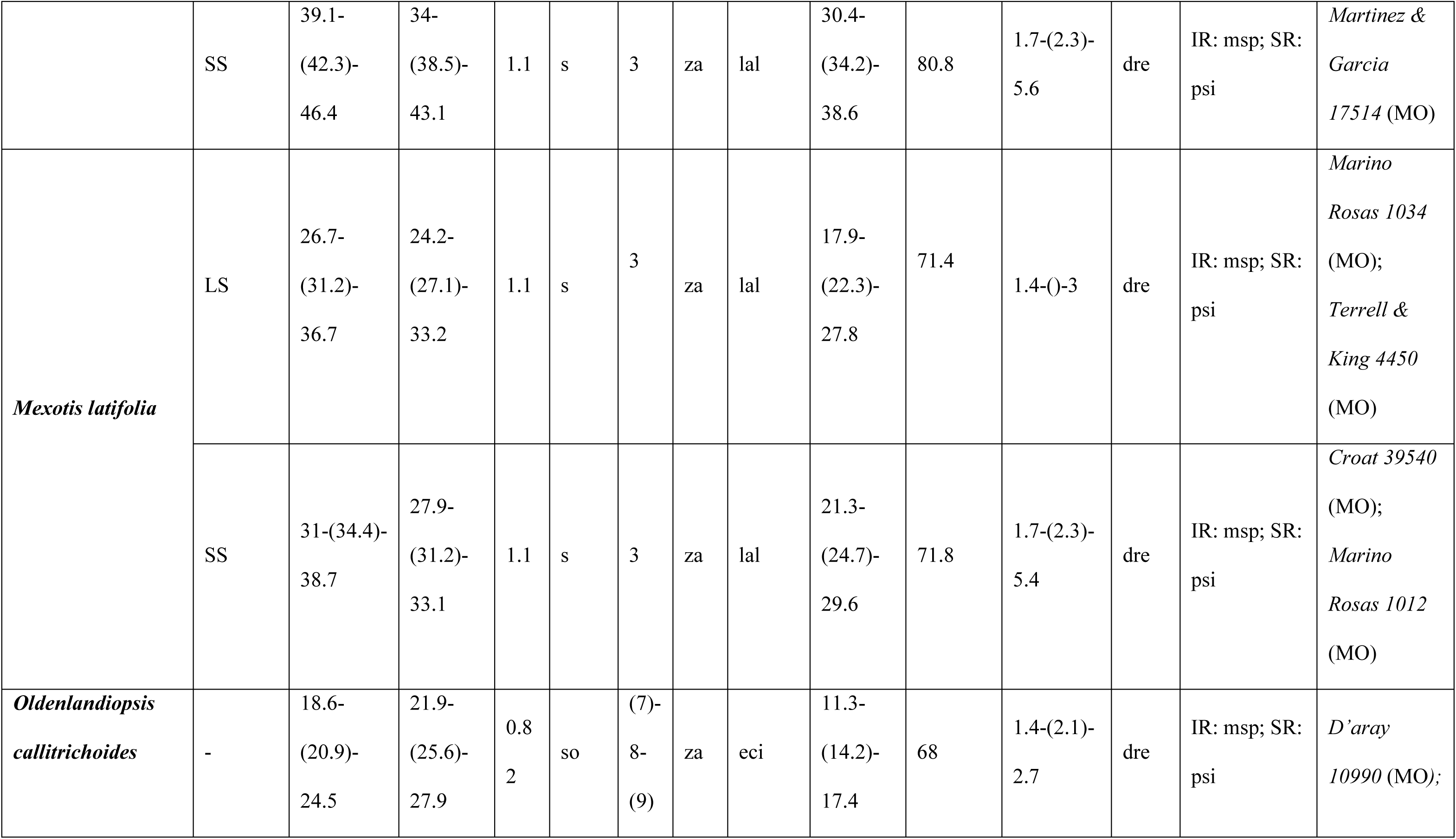
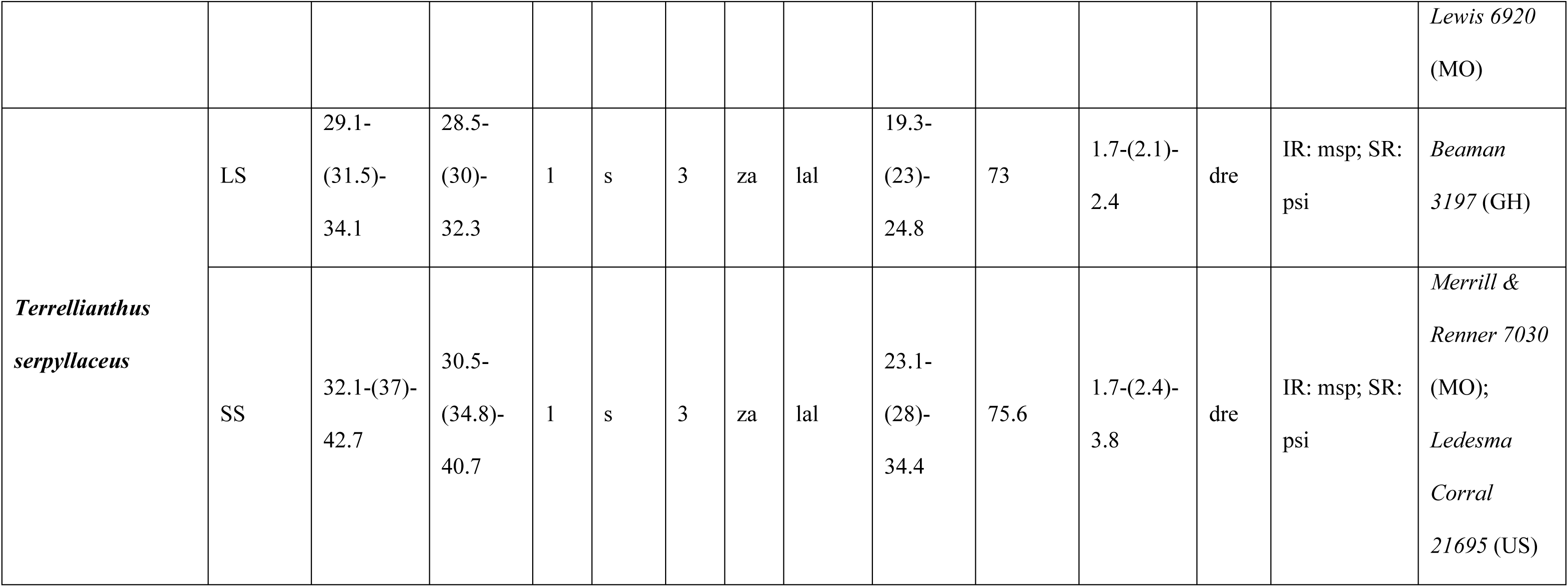
Pollen morphology summary. Abbreviations: dre: double reticulum; E: equatorial diameter; msp: microspines; Endo: endoaperturate; eci: endocingulum; lal: endoaperture lalongate. IR: infrareticulum; Morph: morpho; msp: microspines; LC: length of colpi. LC/P: length of colpi compared with the polar axis; LS: long-styled flower. NC: number of colpi; P: polar axis; Pos: aperture position; psi: psilate; re: reticulate; s: spheroidal; so: suboblate; SR: supra-reticulum; SS: short-styled flower. za: zonoaperturate.

Fruit and seed morphology were examined in four taxa (Fig. 6). *Lucya tetrandra* has capsules wider than long, and cymbiform seeds with a deep ventral depression, thick involute margins, and reticulate-areolate testa (Fig. 6A–D). *Oldenlandia selleana* has an ovoid, 4-lobed capsule, dehiscing loculicidally, and slightly trigonous, angulate seeds, somewhat dorsiventrally compressed, with hilum punctiform, extra-centric, with a reticulate-areolate testa, cell walls slightly undulate anticlinal walls and finely punctate surface (Fig. 6E–H).

**Figure 6.**
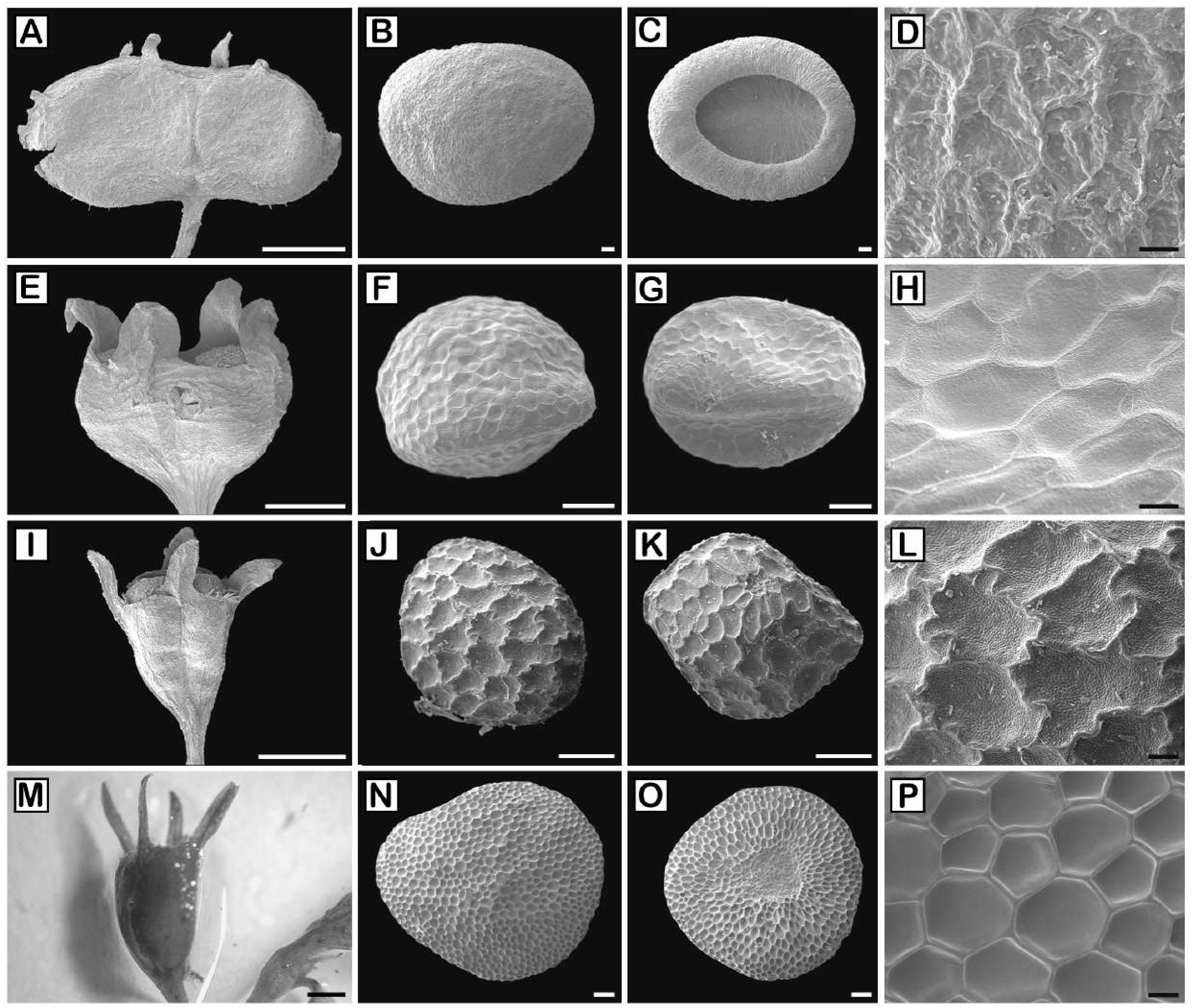
Fruit and seed microcharacters of some Spermacoceae taxa. A–L, N–P. SEM photographs. M. SM photographs. A–D. *Lucya tetrandra*. E–H. *Oldenlandia selleana*. I–L. *Oldenlandiopsis callitrichoides*. M–P. *Terrellianthus serpyllaceus*. A, E, I, M. Fruit. B, F, J, N. Seed, dorsal view. C, G, K, O. Seed, ventral view. D, H, L, P. Detail of exotesta. Scale bars. A, E, I, M. 1 mm. B–C, F–G, J–K, N–O. 100 µm. D, H, L, P. 20 µm. A–D. *Harris 2316* (US). E–H. *Liogier 13024* (NY). I–L. *D’aray 10190* (MO). M–P. *William 22771* (US).

*Oldenlandiopsis callitrichoides* has capsular fruits longer than wide, with a notorious beak, dehiscing loculicidally, and oblate-subglobose seeds, with hilum punctiform, centric, and reticulate-areolate testa bearing sinuous walls, or becoming straight close to the hilum, and finely papillose surface (Fig. 6I–L). *Terrellianthus serpyllaceus* also produces capsules longer than wide, dehiscing loculicidally, with dorsiventrally compressed, suborbicular seeds, and a reticulate-foveolate testa, with thick areoles more or less isodiametric or polygonal (Fig. 6M–P).

## 4. Discussion

The results presented (Figs. 2–3) are broadly consistent with those of previous studies adopting a tribal-level perspective (Kårehed et al. 2008; Groeninckx et al. 2009a; Neupane et al. 2015; Gibbons 2020, among others). Therefore, we focus on those taxa whose phylogenetic position is resolved here for the first time, as well as on clades and relationships not previously reported.

### 4.1. The Dentella-Pentodon clade

The *Dentella*-*Pentodon* clade was resolved in all previous studies, although its placement differs among them (Groeninckx et al. 2009a; Wikström et al. 2013; Gibbons 2020). Here, this clade is resolved as sister to the ALE clade + *Hedyotis*, in accordance with the findings of Kårehed et al. (2008) and Guo et al. (2013). *Dentella* J.R.Forst. & G.Forst. has an Asian-Australian distribution, with *D*. *repens* introduced in Madagascar and Central America; whereas *Pentodon* Hochst. has a geographic range that extends from Africa to the Arabian Peninsula, with *P*. *pentandrus* also introduced to Central and South America (POWO 2026). Both genera share 5-merous flowers, an unusual condition within the tribe, also found in only two other taxa: *Leptoscela* Hook. f., a monospecific, poorly known genus (Nuñez Florentin and Salas 2026), and the recently described *Oldenlandia bahiana* Nuñez Florentin (Nuñez-Florentin et al. 2026), both of which are endemic to Brazil. In both cases, repeated attempts to obtain DNA sequences from the latter two taxa have been unsuccessful. In this sense, further molecular phylogenetic data are needed to confirm the placement of *Leptoscella* and *O*. *bahiana* within Spermacoceae and to determine whether their merosity reflects a phylogenetic affinity with the Dentella-Pentodon clade.

### 4.2. Oldenlandia selleana and the Agathisanthemum–Lelya clade

*Oldenlandia selleana* is a Caribbean taxon endemic to Hispaniola, known only from its original description as a small perennial herb (6–10 cm tall) (Urban 1919). Its phylogenetic position is resolved here for the first time, with the species recovered within the ALE clade as sister to *Lelya* Bremek. and a paraphyletic *Agathisanthemum.* Because Urban (1919) provided no information on fruits or seeds of *O*. *selleana*, SEM examination here carried out striking similarities in seed micromorphology (Fig. 6E–H) with both *Agathisanthemum* and *Lelya* (Bremekamp 1952): angulate seeds with nearly straight to slightly undulate anticlinal walls and a finely punctate surface. The perennial habit with a lignescent base and coriaceous leaves with recurved margins further supports this relationship. Nonetheless, differences in inflorescences, fruit features, and the disjunct distribution between Haiti and Africa remain noteworthy. This disjunction may reflect long-distance dispersal (Janssens et al. 2016), a pattern documented elsewhere in the ALE clade, which is otherwise predominantly African, with the exception of some *Edrastima* species from Asia and the Americas (Neupane et al. 2015).

*Agathisanthemum* is paraphyletic as currently circumscribed, with *Lelya osteocarpa* nested within it with maximum support, an outcome consistently recovered across previous molecular studies (Kårehed et al. 2008; Guo et al. 2013, Groeninckx et al. 2009a; Wikström et al. 2013; Neupane et al. 2015; Gibbons 2020). This result strongly supports combining *L. osteocarpa* under *Agathisanthemum*. Both genera share a distinctive pollen aperture type (“compound ora”), characterized by a distinct endocolpus or endocingulum, a mesoporus surrounded by a costa on the inner grain wall, and a microreticulate sexine with granules on the muri facing the lumina (Lewis 1965). *Lelya* is nonetheless distinguished by its unique thick-walled capsule with a solid beak, as well as differences in dehiscence fruit type and inflorescence. In overall appearance, *Lelya* more closely resembles *O*. *selleana* than *Agathisanthemum*: both are small, compact, low-growing perennial herbs, with stems that root at the base and become somewhat woody, contrasting with the more robust, erect habit of *Agathisanthemum*. However, a full morphological reassessment, including more *Agathisanthemum* species, is needed to corroborate the molecular signal and determine whether both taxa should be formally reduced to *Agathisanthemum*.

### 4.3. The African Oldenlandia oxycoccoides and Edrastima

*Oldenlandia oxycoccoides* was described from East Africa by Bremekamp (1959) under subgenus *Anotidopsis* (Hook. f.) K. Schum. based on its non-beaked capsule, decumbent stems that may root at the nodes, broader leaves, heterostylous flowers, and seed coat cells with undulate walls. Bremekamp (1959) noted the close resemblance between *O. oxycoccoides* and *O. bullockii* Bremek., another African species currently treated as a synonym of *Edrastima goreensis* var. *trichocaula* (Bremek.) Y.D. Zhou. In the present study, the phylogenetic position of *O. oxycoccoides* is demonstrated for the first time, with the species nested within *Edrastima* and recovered as sister to *E. goreensis*. Although a beaked capsule was included in Bremekamp’s (1952) circumscription of *Anotidopsis*, he later broadened the subgenus to accommodate O. *oxycoccoides*, which lacks this character (Bremekamp 1959). Likewise, Nandikar and Bramhadande (2021) reported non-beaked capsules in *O. trinervia* Retz., the nomenclatural type of *Anotidopsis*. Together, these observations indicate that the beaked capsule is not a reliable diagnostic character for *Edrastima*. We therefore emend the diagnosis of *Edrastima* proposed by Neupane et al. (2015) to exclude the beaked capsule and instead define the genus by a combination of characters, including an annual herbaceous habit, terminal and axillary flower clusters, a glabrous corolla tube, subglobose stigmas, and trigonous seeds. The phylogenetic position and morphology of *O. oxycoccoides* support its transfer to *Edrastima* (see Section 4.11 Nomenclature changes).

### 4.4. The relationship between Oldenlandia friesiorum and Neanotis

*Neanotis* W.H.Lewis is a tropical and subtropical Asian genus, characterized by pluriaperturate pollen grains (Lewis 1966). *Oldenlandia friesiorum* is an African species native to Kenya and Tanzania (Bremekamp 1952; POWO 2026), whose close phylogenetic relationship with *N*. *gracilis* is evidenced here for the first time. Comparison of the overall appearance between both taxa based on type specimens indicate that they are similar in the subshrub habit, ovate leaves, and terminal inflorescences. However, since the pollen morphology is a key diagnostic character for *Neanotis*, a detailed palynological study of *O. friesiorum* and related African taxa is forthcoming to determine whether *O*. *friesiorum* should be referred to *Neanotis*.

### 4.5. Dolichocarpa sister to Leptopetalum

*Dolichocarpa* is a recently defined genus based on Australian *Oldenlandia* species (Gibbons 2020). In that work, *Dolichocarpa* was retrieved as sister to the Pacific *Kadua* but with weak support (BS = 57, BPP = 0.74). In the present study, that relationship is not recovered; instead, *D. spermacocoides* is placed with maximum support as sister to a clade comprising all sampled *Leptopetalum* species (BS = 100, BPP = 1). This strongly supported placement suggests a closer evolutionary relationship between *Dolichocarpa* and *Leptopetalum*, and warrants further morphological and molecular investigation to better understand the relationships among them and allied genera of the AAP clade.

### 4.6. The Arcytophyllum-Houstonia complex

This clade is composed of the genera *Arcytophyllum*, *Carterella*, *Houstonia*, *Stenotis*, and *Stenaria*. It is the first time that an analysis has included several representatives per genus, including species of *Stenaria* and *Stenotis*. However, our plastome phylogeny exposes a consistent pattern of morphological homoplasy, with topologies which are incompatible with the current generic circumscriptions proposed by Terrell (1987, 1996, 2001a, b, 2007) and Terrell and Robinson (2010a,b, 2011).

Subclade A comprises *Houstonia* s.str. (*H. caerulea* L., *H. procumbens* (Walter ex J.F.Gmel.) Standl., *H. teretifolia*), all sampled *Stenaria*, and, unexpectedly, *Arcytophyllum fasciculatum*. *Stenaria* was recovered as non-monophyletic with respect to *Houstonia*, corroborating earlier molecular studies that demonstrated *Stenaria* to be nested within the *Houstonia* lineage (Church 2003; Shanks 2015). Terrell (2001a) established *Stenaria* primarily on seed morphology and chromosome number, separating it from *Houstonia* by its non-crateriform ellipsoid seeds with centric punctiform hila and chromosome numbers of x = 9–10, whereas *Houstonia* was characterized by crateriform seeds and the base chromosome numbers x = 6, 7, 8, and 11 forming a descending aneuploid series. Similarly, Terrell and Robinson (2010a) transferred *Houstonia fasciculata* A. Gray (as *Hedyotis intricata* Fosberg) to *Arcytophyllum* following a morphological comparison emphasizing fasciculate branching, condensed woody stems, revolute leaves, and ridged punctiform seed hila shared with some Andean *Arcytophyllum* species. Church (2003) had already suggested a possible affinity of this taxon with the *Houstonia/Stenaria* lineage, although the placement remained unresolved with the data then available. Our phylogeny strongly supports this earlier hypothesis: *A. fasciculatum* is recovered as sister to the Houstonia-Stenaria clade with strong support, whereas the true *Arcytophyllum* (represented here by *A. nitidum*, *A. ericoides*) are placed elsewhere in the complex alongside *Stenotis*, *Carterella*, and *H. prostrata*. Biogeographic patterns reinforce this interpretation, as the native range of *A. fasciculatum* extends from the south-central United States to northeastern Mexico, whereas other *Arcytophyllum* species are distributed primarily in high-elevation regions of Central and South America (Mena 1990).

Furthermore, *A. fasciculatum* possesses a chromosome number of n = 12 (Powell and Leavitt 2011), extending the chromosome diversity observed within the broader *Houstonia–Stenaria* lineage and remaining consistent with the descending aneuploid trend observed in the group. These results indicate that the seed characters emphasized by Terrell are subject to substantial homoplasy and that the morphological features used to place *A. fasciculatum* within *Arcytophyllum* likely represent convergent adaptations rather than indicators of phylogenetic affinity. Similarly, *Stenaria umbratilis* is recovered in a polytomy related to *H. teretifolia,* distant from the remaining *Stenaria* species, which form a separate supported clade. This placement is consistent with morphological differences: unlike the heterostylous flowers typical of *Stenaria* (Terrell 2001a), *S*. *umbratilis* is apparently homostylous, has a creeping, node-rooting habit (vs. non-creeping in other *Stenaria* spp.), and has ovate to broadly elliptic leaves (vs. filiform, linear, narrowly elliptic). A forthcoming study focused on *Houstonia-Stenaria* species complex will further test the relationship between these genera and clarify the placement of *S*. *umbratilis*.

A parallel pattern of non-monophyly was recovered in the second major clade of the complex, which united *Stenotis*, *Carterella alexanderae*, and *Houstonia prostrata* with strong support, sister to Andean *Arcytophyllum* s.str. The present plastome phylogeny is incompatible with the generic limits established by Terrell (1987, 2001b, 2007), who segregated *Stenotis* on the basis of x = 13, colporate type A pollen, and ellipsoid non-crateriform seeds with a centric punctiform hilum; placed *Carterella* in its own monotypic genus based on elongate corollas, laterally compressed angular seeds, and n = 13; and retained *H. prostrata* within *Houstonia* subgenus *Porotis* Terrell based on longitudinally bowed seeds with thickened involute margins, a hilar ridge in a ventral depression, and a uniquely porose testa. As demonstrated above for *Stenaria* and *A. fasciculatum*, the seed characters emphasized by Terrell are subject to substantial homoplasy and cannot reliably delimit genera within this complex. The elongate corollas of *Carterella* (30–50 mm; Terrell 1987) likewise represent a derived floral elongation within the *Stenotis* lineage. Terrell and Robinson (2010b) later acknowledged that *Carterella* was closely related to *Stenotis* based on morphology, cytology, geography, and molecular evidence, although they continued to maintain generic separation primarily because of its highly differentiated floral and seed morphology.

Three independent lines of evidence support recognition of a single lineage. 1) The base chromosome number x = 13, unique within the complex, is documented in both *Stenotis* and *Carterella* (Lewis 1962; Terrell et al. 1986); although the chromosome number of *H*. *prostrata* remains unknown, its phylogenetic placement predicts affinity with this cytological lineage. (2) All three taxa are endemic or near-endemic to Baja California and the Sonoran Desert, with only marginal extensions into the southwestern United States, contrasting with the broad North American distribution of *Houstonia* s.l. (3) according to Lewis (1965) and Terrell et al. (1986) *Stenotis* species and *H*. *prostrata* share 3-colporate pollen grains, with endoapertures lalongate or A ora type as denominated by Lewis. Here, we confirm the same morphological pattern for *C*. *alexanderae*. We therefore propose that *C*. *alexanderae* and *H*. *prostrata* be transferred to an expanded *Stenotis* based on their shared cytological, biogeographic, palynological, and phylogenetic affinities.

### 4.7. The Neotropical Lucya and Oldenlandiopsis, sister to primarily African taxa

The phylogenetic placement of *Lucya* is reported here for the first time, recovered as sister taxon to a clade comprising *Oldenlandiopsis* + *Cordylostigma* + *Oldenlandia* s.s.

Within this clade, *Oldenlandiopsis* is recovered with high support as sister to *Cordylostigma* + *Oldenlandia* s.s., differing from Church (2003), who recovered *Oldenlandiopsis* as sister to *Oldenlandia salzmannii* based on *trnL* data, although without bootstrap support. Both genera are monospecific and restricted to the Neotropics: *Lucya tetrandra* is native to the Caribbean (Puerto Rico, Cuba, Dominican Republic, Haiti, and Jamaica; Terrell 2007), while *Oldenlandiopsis callitrichoides* ranges from southern Mexico through Central America to the Caribbean (Terrell and Lewis 1990).

Both taxa are also morphologically distinct. *Lucya* is perennial with tubers, having homostylous flowers, 6–8 calyx lobed, and cymbiform seeds (Terrell 2007). *Oldenlandiopsis* has a creeping habit and turbinate to obconic, thin-walled, somewhat compressed capsules that dehisce loculicidally before splitting into four narrow segments, with and oblate to obtusely angulate seeds bearing markedly sinuous walls (Terrell and Lewis 1990; Terrell and Robinson 2007). Pollen characters have also been reported to separate both genera from related genera such as *Oldenlandia* and *Houstonia* (Lewis 1966; Terrell and Lewis 1990; Terrell 2007), although no images or detailed palynological descriptions were previously published. We confirm here 6-zonocolporate pollen in *Lucya* and (7-)8(−9)-zonocolporate pollen in *Oldenlandiopsis*; together with *Neanotis*, these are the only genera outside the Spermacoce clade with pluriaperturate (5–12) pollen grains, contrasting sharply with the 3-colporate pollen grains of *Cordylostigma* and *Oldenlandia* s.s. (Lewis 1965; Groeninckx et al. 2010c). *Cordylostima*, although represented by a single species, is consistently recovered as sister to *Oldenlandia* s.s, congruent with previous studies (Kårehed et al. 2008; Groeninckx et al. 2009a; Groeninckx et al. 2010c; Gibbons 2020).

### 4.8. The South American Oldenlandia species

*Oldenlandia dusenii* is a poorly known taxon, except for its treatment in local floras (Delprete et al. 2005), mainly due to its restricted distribution to high-altitude regions in southern Brazil (Paraná, Santa Catarina, and Rio Grande do Sul). Preliminary ongoing morphological studies (Nuñez-Florentin ined.) suggest that this species has distinctive characters in floral, carpological, and palynological morphology with respect to the remaining currently circumscribed South American *Oldenlandia* species (including the adventive *O*. *corymbosa*). The phylogenetic position of this enigmatic taxon is determined for the first time in the present analysis, where it is recovered as closely related to *Manettia cordifolia*. *Manettia* comprises herbs or subshrubs, generally climbers, with seeds that are strongly flattened dorsiventrally and have a noticeable winged margin (Terrell and Robinson 2004; Cordeiro Marinero et al. 2012); whereas *O*. *dusenii* is a creeping herb with ellipsoid, ovoid-outlined, wingless seeds that are not dorsiventrally flattened (Delprete et al. 2005; Nuñez-Florentin ined.) Despite these differences, they seem to share the leaf shapes, oval to cordate leaves, and septicidal dehiscent fruits. In terms of phylogeny, Firens da Silveira (2015) conducted the most comprehensive molecular phylogenetic study of *Manettia* and confirmed its monophyly, although the sister group of the genus could not be determined.

*Bouvardia* was proposed as a probable sister group, a relationship not recovered or supported in the present analysis. Given the results of Firens da Silveira (2015) and those obtained here, it is clear that further morphological and phylogenetic studies including a broader sampling of *Manettia* species are needed to elucidate the intergeneric relationship between *Manettia* and *O*. *dusenii*. Nevertheless, there is no doubt that *O*. *dusenii* represents a distinct lineage from the other native South American (*O*. *salzmannii* lineage) or adventive to S. America (*O*. *corymbosa* lineage) *Oldenlandia* species.

In the present analysis, the widespread *O*. *salzmannii* is recovered as sister taxa to the Spermacoce clade. This is congruent with Kårehed et al. (2008), although there *O. salzmannii* formed a well-supported clade with *O*. *tenuis* K.Schum. A lot of effort to obtain plastome information from *O*. *tenuis* was made, but without success. *Oldenlandia salzmannii* is characterized as a creeping marsh herb with ovate leaves and heterostylous flowers. It is widely distributed throughout South America and introduced in Florida, United States (Fosberg and Terrell 1985; Nuñez-Florentin et al. 2016). In contrast *O*. *tenuis* is distinguished by its slender habit, linear leaves, homostylous flowers, and is distributed in northeastern Brazil and Venezuela (Steyermark 1988). Both species share distinctive palynological features 4–5-apertured pollen grains with an endocingulum as the endoaperture (Nuñez Florentin, ined.). These traits distinguish them from *Oldenlandia* s.s. species, a predominantly African lineage characterized by 3–(4)-colporate pollen grains with an lalongate endoaperture (Lewis 1964; Neupane et al. 2015). Due to these micromorphological differences, geographical distribution, and phylogenetic evidence, the South American species currently assigned to *Oldenlandia* likely represent a lineage distinct from *Oldenlandia* s.s.. Additional studies integrating morphological and phylogenetic data will be necessary to determine the relationships of these native southamerican species to one another and to related genera.

### 4.9. The Bouvardia complex and related taxa

The Bouvardia clade is resolved as a sister group to the Manettia clade, although with weak support. This clade is strictly American and includes several taxa whose phylogenetic position is reported here for the first time, such as *Martensianthus galleotti* and *Mexotis latifolia*. The analysis placed them within the *Bouvardia* clade, a not surprising position, since both taxa were taxonomically related to *Bouvardia* (Terrell and Robinson 2009; Borhidi and Lozada-Pérez 2010, 2011). Both, *Martensianthus* and *Mexotis* are mainly distributed in Mexico and the southern United States of America and share certain morphological characteristics with *Bouvardia*, such as flattened, often winged seeds and terminal inflorescences (Borhidi and Lozada-Pérez 2010; Terrell and Robinson 2009). We confirm here the presence of 3-colporate pollen grains with a lalongate endoaperture for both taxa, a palynological feature that links them to *Bouvardia* (Arreguín-Sánchez et al. 1995). The species currently included in *Mexotis* were separated from a group of American *Hedyotis* species (Terrell and Robinson 2009). Borhidi and Lozada-Pérez (2010, 2011) excluded *M*. *galeottii* from *Mexotis* and transferred it, along with four other *Bouvardia* species, to the newly established genus *Martensianthus*. This genus was differentiated from *Bouvardia* primarily by corolla characteristics (funnel-shaped corolla, white, 3–10 mm long, with densely pubescent lobes vs. hypocrateriform corolla, varied in color, 7–45 mm long, generally glabrous internally). The phylogenetic analyses presented here confirm for the first time that *Bouvardia*, *Mexotis*, and *Martensianthus* form a closely related lineage, supporting their recent taxonomic association. However, *Terrellianthus serpyllaceus* is recovered as the sister to the genus *Bouvardia*.

*Terrellianthus* is a monospecific genus from Mexico, whose phylogenetic position has been widely debated. Originally described as *Hedyotis serpyllacea* Schltdl., it was transferred by Terrell (1999) to *Arcytophyllum*, based on habit and seed morphology. Andersson et al. (2002) established a re-circumscribed *Arcytophyllum* based on molecular evidence (*rps16* intron), finding that *A*. *serpyllaceum* does not fall within the rest of the genus, but is closely related to *Bouvardia* and *Manettia*. Studies by Kårehed et al. (2008) and Groeninckx et al. (2009b) obtained similar results, supporting the circumscription of Andersson et al. (2002).

However, Terrell and Robinson (2004) rejected Andersson’s re-circumscription, noting that they were unable to confirm the identity of the sampled material because the voucher specimen (Stafford et al. 203, MO) was not found in the herbarium where it was purportedly deposited. Years later, Borhidi (2012) proposed a solution to this conflict by creating the new genus *Terrellianthus* for *A*. *serpyllaceum*. Based on his observations, he considered it closely related to the Bouvardia-Martensianthus complex due to the similarities in its stipules, flowers, and loculicidally dehiscent fruits. In the present analysis, the position of *T*. *serpyllaceus* within the Bouvardia clade is confirmed using material different from that analyzed by Andersson et al. (2002). Nevertheless, future studies incorporating a diversity of *Bouvardia* species will be necessary to determine whether the current generic limits should be expanded to include these related taxa.

### 4.10. The Spermacoce clade

In the *Spermacoce* clade, the backbone topology and major clades are congruent with those recovered in the recent studies (Nuñez Florentin et al. 2024; Florentín et al. 2025).

Although the Galiantheoid and the Homostyloid clades were originally defined by Florentín et al. (2025), both are recovered here with strong support. In previous studies, *Galianthe* (Neotropical) was recovered as monophyletic within the Galiantheoid clade, with Schwendenera (endemic to Brazil) as its sister genus (Florentín et al. 2025). In contrast, our analyses recover *Galianthe*, as currently circumscribed, as polyphyletic. *Galianthe vaginata* forms a strongly supported clade with *Schwendenera tetrapyxi*s, whereas the remaining *Galianthe* species included in our analyses form a strongly supported clade sister to *Carajasia cangae* (endemic to Brazil), although support for this sister relationship is weak. These three genera share distinctive palynological features, including pollen grains with long colpi, a double reticulum, and the absence of orbicules (Salas et al. 2015a; Nuñez Florentin et al. 2023). Nevertheless, *Carajasia* was distinguished from *Galianthe* by its uniflorous inflorescences and homostylous flowers (Salas et al. 2015a). However, during the last decades, the traditional concept of *Galianthe*, having only heterostylous flowers, has been expanded to include new homostylous species, such as *G*. *holmneielsenii* Florentín & R.M.Salas, *G*. *palustris* (Cham. & Schltdl.) Cabaña Fader & E.L.Cabral, *G*. *spicata* (Miq.) Cabaña Fader & Dessein, and *G*. *vasquezii* R.M.Salas & Florentín (Florentín et al. 2017), weakening the diagnostic value of *Carajasia*. Despite the current phylogenetic signal to merge *Carajasia*, *Galianthe*, and *Schewendenera*, we considered that a comprehensive sampling of the subclades identified in Florentín et al. (2025), within Galianthe as Cymoid, Ebeloid, Galianthe s.l., and Galianthe s.s. clades are needed before applying such a taxonomic decision.

*Spermacoce* remains polyphyletic, consistent with previous studies (Salas et al. 2015a; Miguel et al. 2018; Nuñez-Florentin et al. 2024). Five Australasian species — *S. auriculata* F.Muell.*, S. occidentalis* Harwood*, S. papuana* F.Muell.*, S. pogostoma* Benth., and *S. tectanthera* Harwood — are sequenced and included in a phylogenetic analysis for the first time. They are recovered as sister to the American S*. eryngioides* (Cham. & Schltdl.) Kuntze and *S. tenuior* L., together forming a strongly supported *Spermacoce* s.s. lineage. The African species, represented here only by *S. dibrachiata*, is recovered in a distinct clade sister to *Planaltina lanigera* and *Tessiera lithospermoides*, genera endemic to Brazil and Mexico, respectively (Salas and Cabral 2010a,c). This study also provides the first phylogenetic evidence for the phylogenetic placement of *Planaltina* and *Tessiera*. The later taxa, *S. dibrachiata*, and *Hexasepalum teres* (Walter) J.H.Kirkbr. share large pollen grains (40-60 μm) with short colpi (Dessein et al. 2002; Salas and Cabral 2010a,c). A comprehensive comparative morphological and phylogenetic study of African, American, Asian, and Australian Spermacoce species, undertaken jointly by multiple research groups, is needed to clarify their interrelationships and their relationships with genera such as *Hexasepalum*, *Planaltina*, and *Tessiera*, among others.

*Borreria*, as currently circumscribed, is a Neotropical genus (Miguel et al. 2022; Souza et al. 2022). The Borreria-Spermacoce complex is perhaps the most contentious taxonomic debate in the history of the tribe Spermacoceae, with numerous studies presenting opposing views (e.g., Dessein 2003; Delprete et al. 2005; Cabral et al. 2012; Miguel et al. 2018). Despite considerable effort, we were unable to obtain reliable plastome data for *Borreria* species, which are therefore excluded from the present analyses, and should be a focus of future study.

### 4. 11. Nomenclature changes

***Edrastima oxycoccoides*** (Bremek.) Nuñez-Florentin & Neupane, **comb. nov**. ≡ *Oldenlandia oxycoccoides* Bremek. in Kew Bull. 13: 382. 1959. – Holotype: Tanzania, Southern Highlands Province, Iringa District, Ngwazi, *Carmichael 359* (EA); Isotype: K [K000316343] ***Stenotis alexanderae*** (A.M. Carter) Nuñez-Florentin & Neupane, **comb. nov.** ≡ *Bouvardia alexanderae* A.M. Carter in Madroño 13(4): 142–144, f. 1–2. 1955. ≡ *Hedyotis alexanderae* (A.M. Carter) W.H. Lewis in Ann. Missouri Bot. Gard. 55(1): 31. 1968. ≡ *Cartelera alexanderae* (A.M. Carter) Terrell in Brittonia 39(2): 250. 1987. – Holotype: Mexico: Baja California Sur: Arroyo del Salto, E of La Paz, 24°12’N, 110°07’.5’W, 30 Mar 1949, *Carter 2577* (UC), [UC-985926]; Isotypes: F [F0068554F], MEXU [MEXU00049905], US [US00137789].

***Stenotis prostrata*** (Brandegee) Nuñez-Florentin & Neupane, **comb. nov.** ≡ *Houstonia prostrata* Brandegee in Zoë 5(6–8): 105. 1901. ≡ *Hedyotis vegrandis* W.H. Lewis in Rhodora 63 (752): 222. 1961. ≡*Houstonia prostrata* var. *prostrata* Brandegee in Shreve & Wiggins, Veg. Fl. Sonoran Desert 2:1399.1964.– Lectotype: Mexico: Baja California: La Palma, Cape Region, 25 Sept 1899, Brandegee s.n. (UC), [UC102445]; Isolectotypes: GH [GH00096790], NY [NY00131925], US [US00137475; US00137476].

= *Houstonia parvula* Brandegee in Zoë 5(10C): 221. 1905.≡ *Hedyotis sinaloae* W.H. Lewis Rhodora 63(752): 222. 1961. *Houstonia prostrata* var. *parvula* (Brandegee) Wiggins, in Shreve & Wiggins, Veg. Fl. Sonoran Desert 2:1399.1964. Lectotype: Mexico: Sinaloa, gravel deposits of Tamazula River near Culiacan, 12 Oct 1904, *Brandegee s.n.* UC [UC102454]; Isolectotypes: E [E00505229], GH [GH00096789, GH00096788], NY [NY00131920], US [US00137471, US00763549].

### 4.12. General Survey of tribe Spermacoceae

Table 2 presents an updated list of all genera currently recognised in the tribe Spermacoceae, together with their geographical distribution and key bibliographic references. Based on recent studies (Verstraete et al. 2026), and mainly on the phylogenetic and taxonomic results presented above, 82 genera are here recognised for the tribe, of which at least seven are tentatively included pending further resolution of their affinities. Of the total genera, 36 are exclusive to the Americas (43.95%), predominantly in the Neotropics; 21 are restricted to Africa (25.6%); 8 exclusive to Asia (9.7%); 3 to Australia and Pacific Islands (3.6%); 9 are considered as Paleotropical in distribution (ca. 10%); and 5 are considered with a Pantropical distribution (4.8%).

**Table 2.**
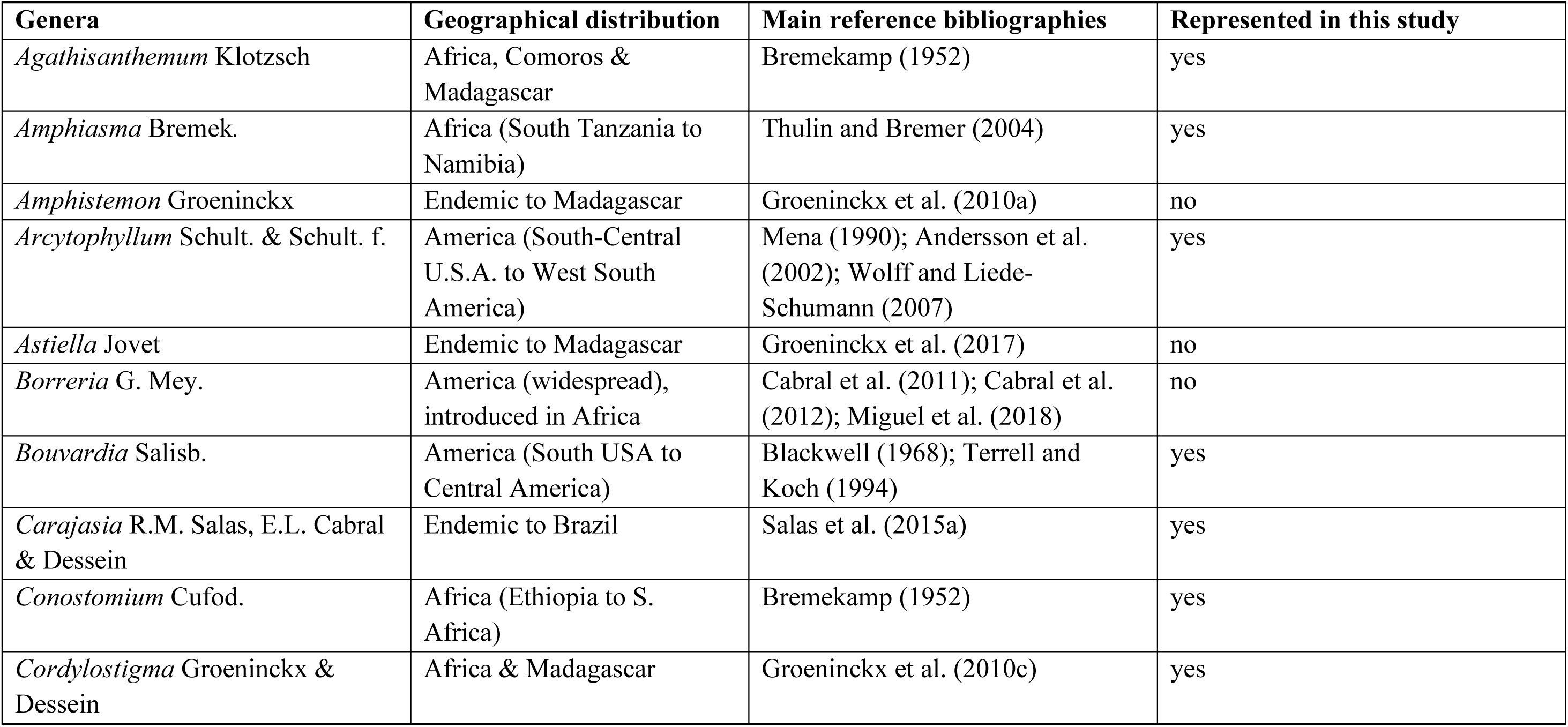
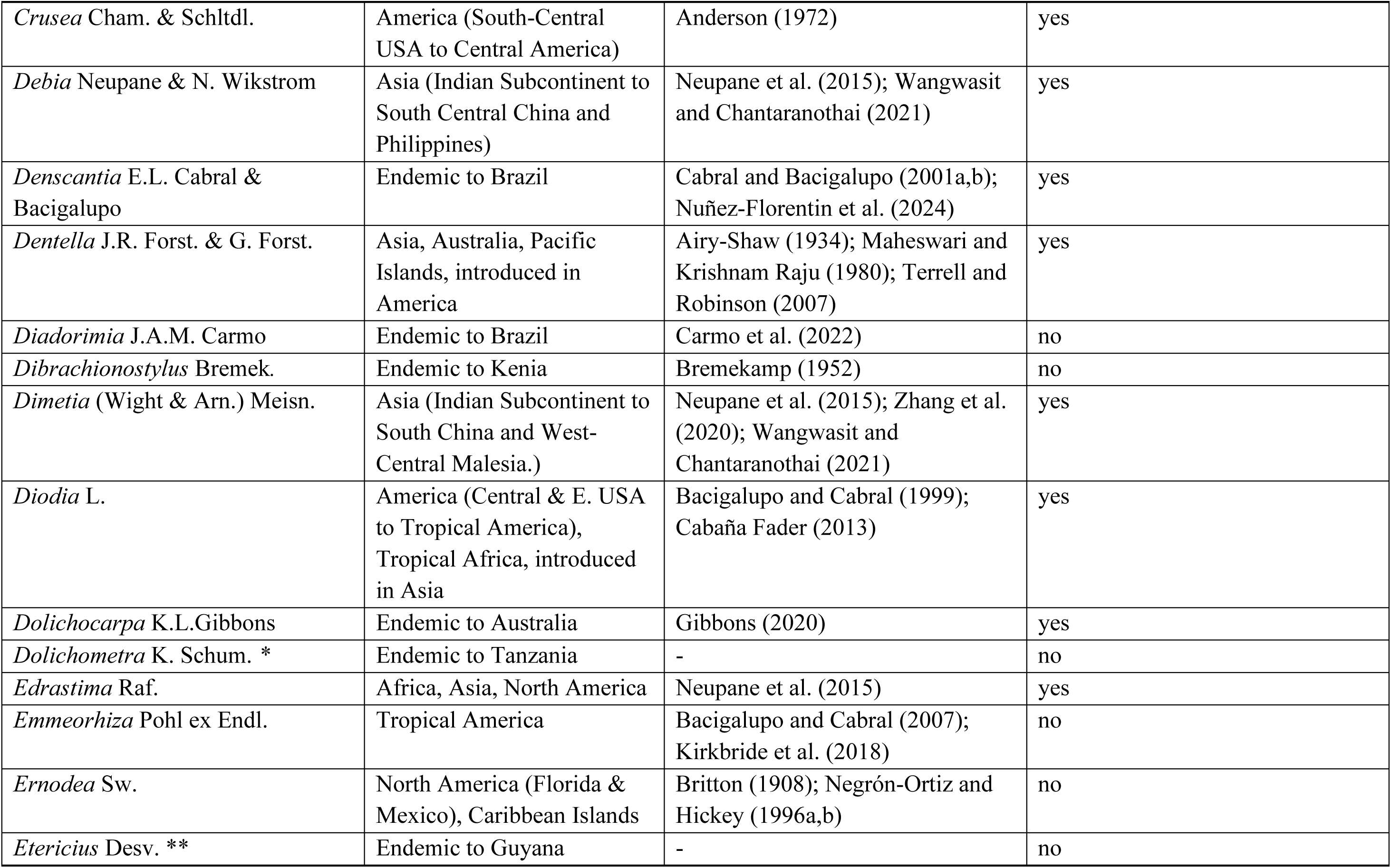
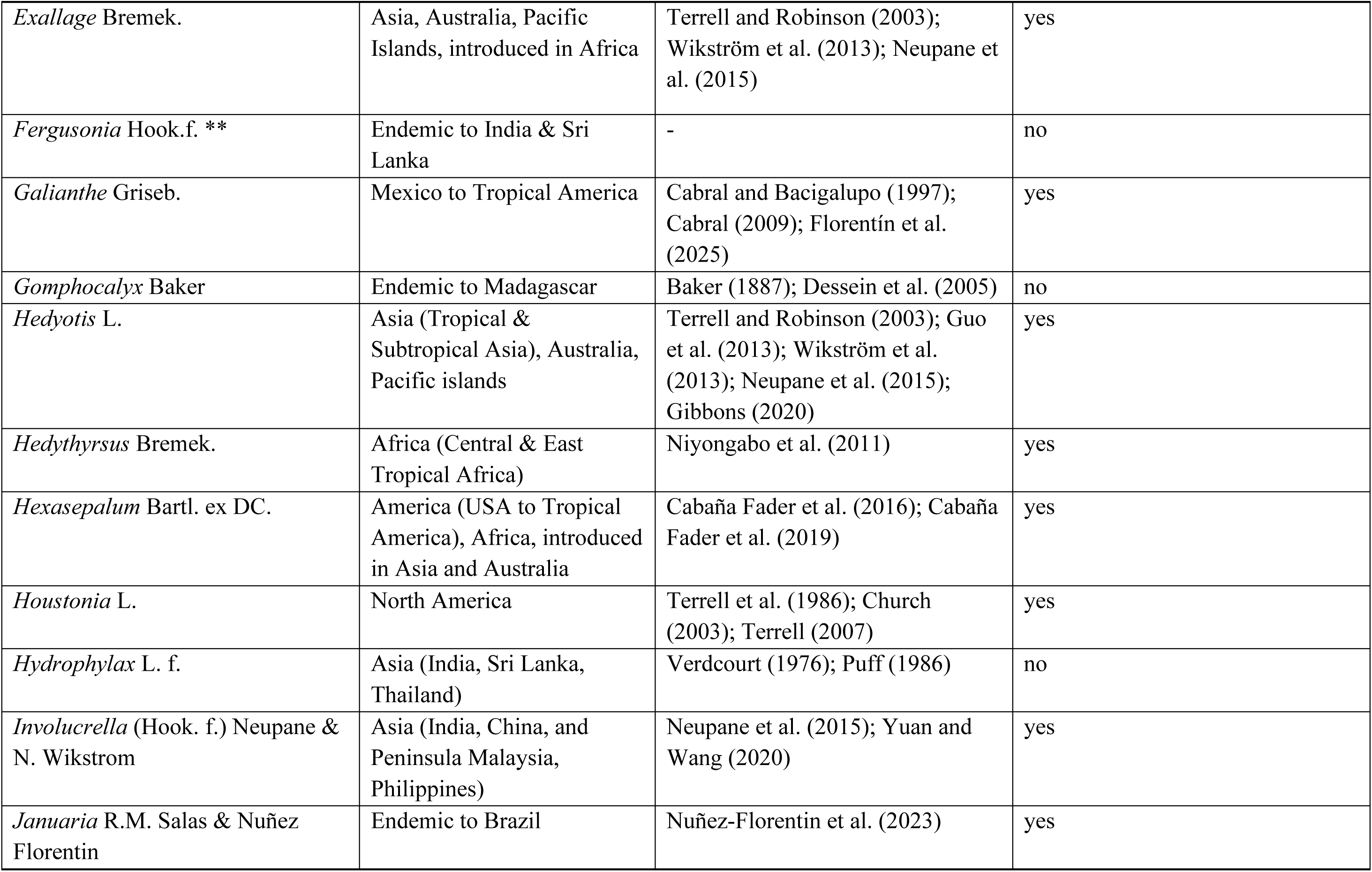
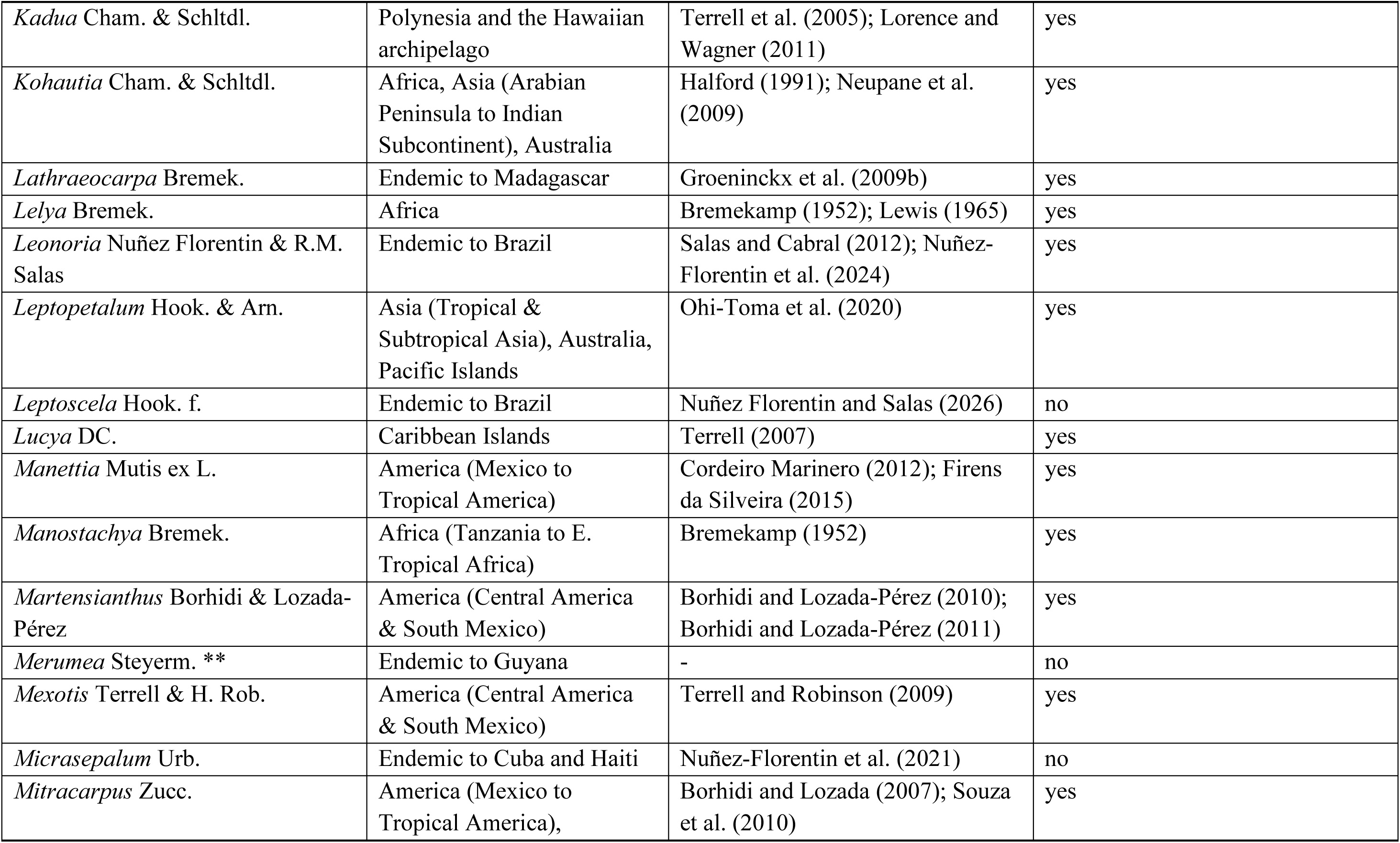
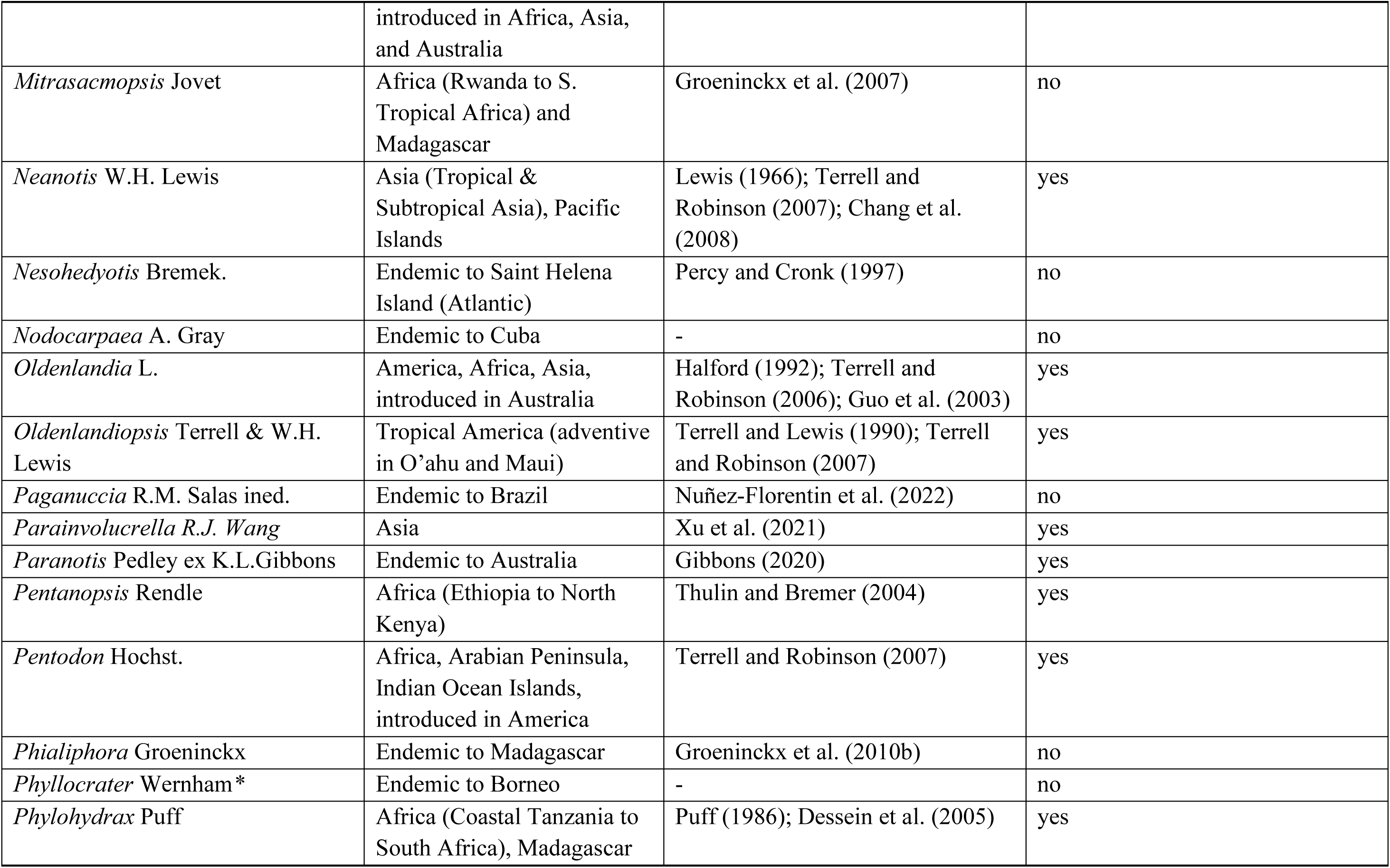
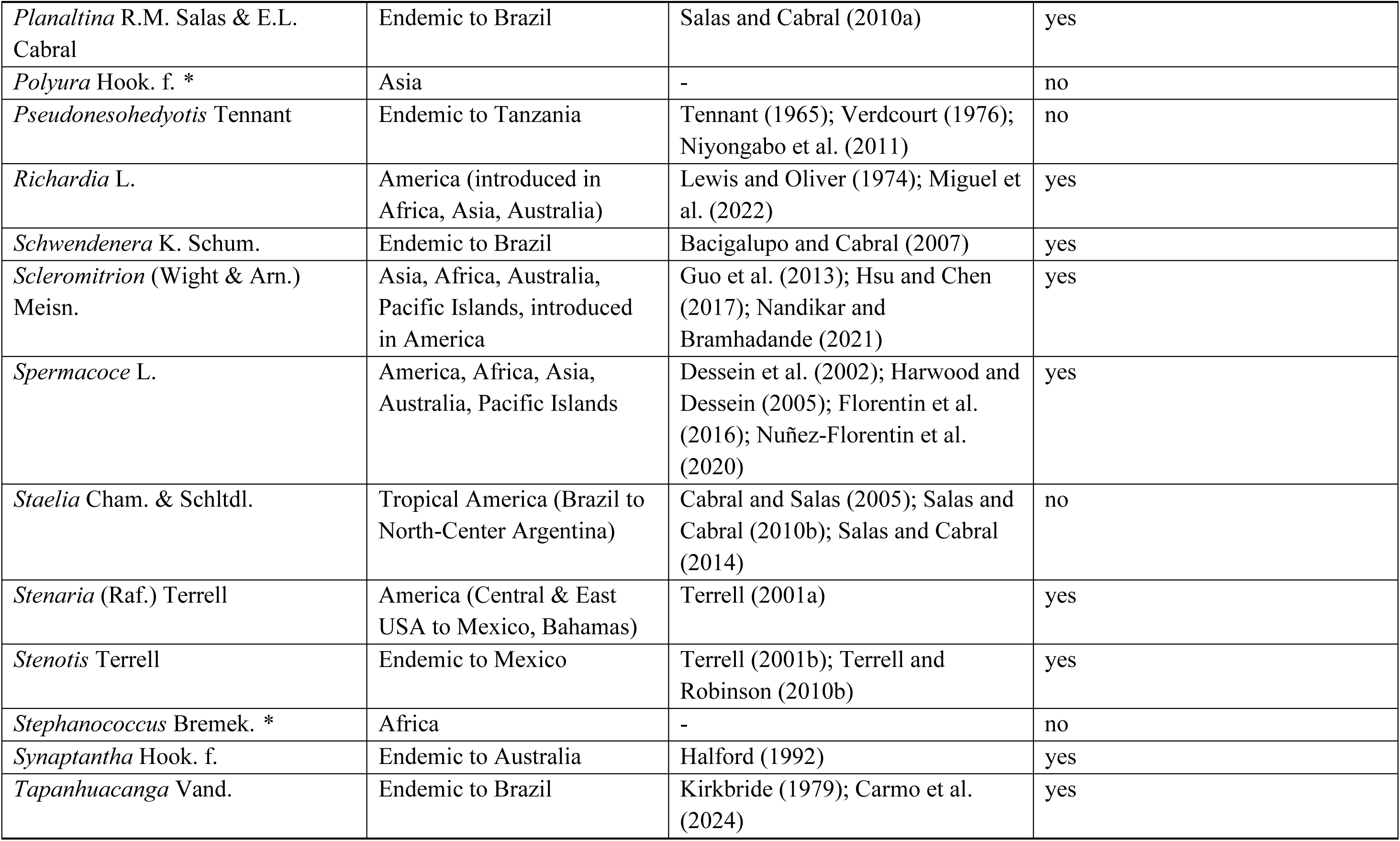
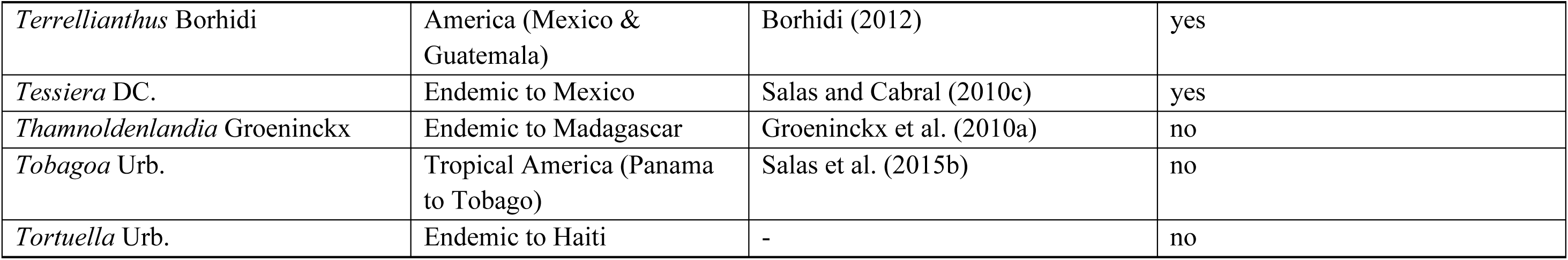
List of all genera currently belonging to the tribe Spermacoceae, including their geographical distribution (based on specific bibliography corroborated with POWO 2026), and main reference bibliographies. (*) Genera listed by Groeninckx et al. (2009a), (**) Genera listed by Razafimandimbison & Rydin (2004); in both cases, those genera lack a taxonomic study or are not included in any phylogenetic study confirming their position in the tribe, then considered tentatively included.

### 4.13. Key to the recognised genera of the tribe Spermacoceae

Below is a summary of all the genera recognized in the tribe Spermacoceae, with the exception of the following taxa, whose inclusion in the key was not possible due to the scarce information available for them: *Dolichometra*, *Etericius*, *Fergusonia*, *Merumea*, *Phyllocrater*, *Polyura*, *Sacosperma*, *Stephanococcus*.

1. Fruit with one seed per locule 2

1’. Fruits with multi-seeded locules (except *Astiella delicatula* Jovet)…36

2. Ovules inserted at the base of the septum (except *Richardia*)…3

2’. Ovules inserted in the middle or just below the middle of the septum 5

3. Calyx 8-lobed, calyx tube very reduced; endemic to Madagascar…4

3’. Calyx 4-lobed, calyx tube well developed; distributed on the east coast of Africa (from Mozambique to South Africa) and Madagascar…***Phylohydrax***

4. Subshrub; stigma 3–4-lobed; ovary 3–4-locular; fruit fleshy, indehiscent ***Lathraeocarpa***

4’. Herbaceous, prostrate or decumbent, sometimes forming dense clumps; stigma bifid; ovary 2-locular; fruit dry, indehiscent…***Gomphocalyx***

5. Inflorescences lax, thyrsoid or pleiotyrsoid, with partial multiflorous inflorescences glomeriform or subglomeriform. 6

5’. Inflorescences mostly compressed; flowers in glomerules, axillary or pseudoaxillary, pauciflorous to multiflorous, or 1–2 flowers…9

6. Plants erect, rarely supporting or climbing; flowers heterostylous, or rarely homostylous; seeds with persistent or deciduous strophiole ***Galianthe***

6’. Plants climbing; flowers always homostylous; seeds with persistent strophiole **7**

7. Fruits dry and indehiscent, or lately separated at the apex into two indehiscent carpels; seeds not winged or with a small winged margin. ***Paganuccia***

7’. Capsule with septicidal dehiscence in two valves; winged seeds; wing derived from the strophiole exceeding the base and apex of the seed or as an extension of the thinned exotesta 8

8. Fruits with the apical portion of the carpels exceeding the pericarp and forming a “beak” or “rostrum”; strophiole developed and exceeding the length of the seed apically and basally; pedicellate flowers, pedicel 1.7–5.5 mm long…***Emmeorhiza***

8’. Fruits without an apical portion exceeding the pericarp; strophiole as long as the seed; flowers short-pedicellate, pedicel 0.5–1.2 mm long…***Denscantia***

9. Calyx tube and lobes undeveloped or developed but fragile and prematurely deciduous 10

9’. Calyx tube and lobes well developed or reduced…11

10. Subshrub erect; flowers 4-merous; corolla 1.3–1.8 cm long…***Micrasepalum***

10’. Herbaceous, rooting at nodes; flowers 3-merous; corolla 0.4–0.5 mm long..***Nodocarpaea***

11. Fruits fleshy, dry indehiscent or schizocarpic 12

11’. Capsules with longitudinal, circumcisely-transverse, or longitudinally-oblique dehiscence 24

12. Heterostylous flowers…13

12’. Homostylous flowers…15

13. Bifid stigma; indehiscent fruits…14

13’. 4-fid stigma; schizocarp fruits separating into 3–4 mericarps…***Schwendenera***

14. Leaves linear or narrowly elliptical, 5–10 × 0.3 mm, leathery when dry; fruits fleshy reddish-orange fruits; endemic to Haiti. ***Tortuella***

14’. Leaves ovate or elliptical, 40–110 × 25–30 mm long, papery when dry; fruit dry, brown; distributed in Panama, Tobago, northern Venezuela. ***Tobagoa***

15. Ovary 3–4-carpellate; stigma 3-lobed, 4-lobed, rarely 2-lobed…***Richardia***

15’. Ovary 2-carpellate; stigma bifid, 2-lobed, or obscurely 2–4-lobed…16

16. Dry fruits schizocarpous or indehiscent; (lately separated at the apex into two indehiscent carpels); seeds not ruminated [except *Hexasepalum gardneri* (K. Schum.) J.H. Kirkbr. & Delprete]…17

16’. Fruits fleshy, indehiscent, reddish-orange; seeds ruminated. ***Ernodea***

17. Inflorescences pauciflorous (1–5 flowers per node)…18

17’. Pluriflorous inflorescences (40–100 flowers per node)…22

18. Rocky herbs, with reddish stems and leaves; calyx lobes 0.15–0.3 mm long; fruits schizocarps, separating into two mericarps leaving a basal carpophore ***Carajasia***

18’. Herbs or subshrubs, generally psammophilous or growing in moist soil, rarely rupicolous, with greenish stems and leaves; calyx lobes 0.6–8 mm long; fruits schizocarps separating into two mericarps without basal carpophore, or indehiscent fruits (late separated at the apex into two indehiscent carpels). 19

19. Fruits obconic, 15–20 mm long; often asymmetrical, curved longitudinally due to one of the ovules being aborted; endemic to Asia (Andaman Islands, India, Sri Lanka, and Thailand)…***Hydrophylax***

19’. Fruits sub-spherical, ovoid, slightly obconic, 2–10(−12.5) mm long; both ovules well developed; native to America and African coasts 20

20. Growing in moist soils, edge of water bodies; corolla hypocrateriform, corolla tube internally glabrous, corolla lobes internally pubescent; stigma bifid…***Diodia*** s.s.

20’. Growing mainly in sandy, non-marshy soils; funnel-shaped corolla, lower part of the corolla tube internally with a ring of trichomes; stigma 2-lobed or obscurely 2–4-lobed…21

21. Ventral side of mericarps with two depressions; embryo curved; pollen grains with spines and granules…***Hexasepalum*** s.s.

21’. Ventral surface of mericarps flat (without depressions); embryo straight; pollen grains only with microspines…***Hexasepalum*** s.l.

22. Schizocarp fruits separating into two mericarps, leaving a basal carpophore; calyx deciduous, rarely persistent on the fruit; corolla 3–49 mm long, reddish, pink, lilac, violet, blue or white but with coloured lobes at the apex ***Crusea***

22’. Dry indehiscent fruits, late separated at the apex into two mericarps, without leaving a basal carpophore; calyx persistent in the mature fruit; corolla 0.9–4 mm long, corolla white…23

23. Inflorescences axillary glomerules (bilateral); stamens and style exserted; corolla internally with a ring of trichomes in the middle of the tube, lobes glabrous; 1(−2) seeds per capsule ***Borreria*** p.p.

23’. Inflorescences glomerules pseudoaxillary (unilateral), rarely terminal or 1–2 axillary glomerules [e.g. *Spermacoce decipiens* (K. Schum.) Kuntze]; stamens and style included; corolla internally with a ring of trichomes in the lower half of the corolla lobes or in the corolla tube; two seeds per capsule ***Spermacoce*** p.p.

24. Heterostylous flowers; always bifid style 25

24’. Homostylous flowers; style variable: capitate, capitate-bilobate, bilobate, bifid 26

25. Stems cespitose, with a well-developed, woody subterranean system; stipules triangular, with the margins bearing 4 small lobes; capsule dehiscence longitudinal-oblique, the two valves forming one single caducous diaspore ***Diadorimia***

25’. Stems erect, rarely decumbent, with a taproot system; stipules fimbriate; inflorescences in glomerules or pauciflorous cymes; capsule dehiscence septifragal, valves persistent from which the seeds are shed after dehiscence ***Tapanhuacanga*** p.p.

26. Fruits with circumscissile or longitudinally-oblique dehiscence; calyx 2–4-lobed, when 4-lobed, then with 2 lobes markedly smaller than the other two. 27

26’. Fruits with longitudinal dehiscence; calyx 2–4-lobed, all lobes of equal length…28

27. Fruits with circumcisile dehiscence; after dehiscence, the fruit separates into two parts, the upper part in the shape of a ‘mitre’ and comprising the upper portion of the carpels and persistent lobes, and the lower part formed by the basal part of the carpels and the septum; calyx 4-lobed, with 2 longer lobes and 2 shorter lobes, rarely exceptions; corolla mostly hypocrateriform…***Mitracarpus***

27’. Fruits with longitudinal-oblique dehiscence; after dehiscence, the fruit separates into three parts, two valves and a basal portion formed by the basal portion of the carpels, intercarpel septum remains complete, erect and persistent in the plant; calyx 2-lobed (rarely 3–4), with lobes equal in length; corolla funnel-shaped…***Staelia*** p.p.

28. Capsule that separates into three parts, two deciduous valves and a persistent intercarpelal septum 29

28’. Capsule whose dehiscence occurs in two dehiscent valves, or one dehiscent valve and the other indehiscent; both valves remain together and persistent in the plant, with or without an intercarpel septum between 32

29. Capsule strongly compressed laterally; flowers with style and stamens included…***Tapanhuacanga*** p.p.

29’. Capsule obovoid or subglobose; flowers with style and stamens exserted…30

30. Calyx 2-lobed; stigma bifid; pollen grains small (P=25.7; E=25.3 μm), with long colpi; ventral surface of seeds without ruminations…***Staelia catechosperma*** K. Schum.

[previously *Anthospermopsis* (K. Schum.) J.H. Kirkbr.]

30’. Calyx 4–7-lobed; stigma bilobed or obscurely 2–4-lobed; pollen grains large (P=42–75; E=50–78.4 μm), with short colpi; ventral surface of seeds ruminated…31

31. Stems scabrous, pubescent, glabrescent or glabrous; fruits with membranous septum with impressions of the seeds; seeds ruminated on both sides; pollen grains 12–14-colporate, exine with perforations without thickenings ***Tessiera***

31’. Stems woolly, hirsute or hispid; fruits with leathery septum without impressions of the seeds; seeds with invaginations only on the ventral side [except *Planaltina capitata* (K.

Schum.) R.M. Salas & E.L. Cabral]; pollen grains 10–11(−13)-colporate, exine with thick-edged perforations…***Planaltina***

32. Capsule compressed laterally, with a septum, persistent, erect, parallel to the valves…33

32’. Capsules obconic, turbinate, obovoid, not compressed, without membranous septum parallel to the valves…34

33. Flowering branches determinate or indeterminate; inflorescences in 1-flowered cymes or terminal glomerules, rarely pauciflorous cymes; calyx 2-lobate; flowers with style and stamens included; style bilobate, rarely capitate; pollen grains 4–7 zonocolporate, with microspines mostly concentrated along each side of the colpi ***Tapanhuacanga*** p.p.

33’. Flowering branches always indeterminate; inflorescences with axillary glomerules; calyx 4-lobate; flowers with style and stamens exerted; style bifid; pollen grains 8–9 zonocolporate, with microspines and granules uniformly distributed in the exine surface ***Leonoria***

34. Pollen with a reticulate simple exine; longitudinal-transverse dehiscence, with one indehiscent carpel and one dehiscent valve that separates from the pedicel…***Januaria***

34’. Pollen with an eutectate, perforate, rarely microreticulate exine; longitudinal dehiscence, with both carpels dehiscent or both indehiscent, or one carpel indehiscent and one valve dehiscent, in all cases both valves remain together and persist on the pedicel…35

35. Inflorescences in axillary glomerules (bilateral); stamens and style exserted; stamens inserted in the interlobular sinuses of the corolla ***Borreria*** p.p.

35’. Inflorescences in pseudoaxillary glomerules (unilateral), rarely terminal and 1–2 axillary; stamens and style included; stamens inserted in the middle or base of the corolla tube, or in the upper half of the corolla tube…***Spermacoce*** p.p

36. Plants dioecious; small trees up to 7 m tall; flowers unisexual; stipules connate into a tubular sheath; endemic to St. Helena ***Nesohedyotis***

36’. Plants no dioecious; herbs, subshrubs, shrubs, rarely small trees; flowers perfect, homostylous or heterostylous: stipules fimbriate, lobate, rarely tubular; distributed in Australia, Africa, Asia, Polynesia or the Americas, never restricted to St. Helena alone…37

37. Flowers always 5-merous…38

37’. Flowers 4-merous…40

38. Creeping, prostrate herbs; flowers homostylous; dry indehiscent fruit ***Dentella***

38’. Erect, decumbent, or voluble herbs or shrubs; flowers heterostylous; dry dehiscent fruit………………………………..39

39. Inflorescences in cincinnus, axillary; capsule obovoid, with septicidal dehiscence; seeds 0.7–1 mm long, ovate to suborbicular ***Leptoscela***

39’. Inflorescences not in cincinnus, terminal or axillary; capsules oblong, with loculicidal dehiscence; seeds 0.3–0.5 mm long, trigonous…***Pentodon***

40. Indehiscent fruits…41

40’. Dehiscent fruits…43

41. Terminal inflorescences, subtended by 4 bract-like leaves; dry indehiscent fruits…Parainvolucrella

41’. Axillary inflorescences, or if terminal never subtended by 4 bract-like leaves; fleshy or cartilaginous fruits…42

42. Inflorescence always axillary cymes; seeds trigonous…***Exallage***

42’. Inflorescence terminal, paniculate; seeds dorsiventral compressed, thickened, irregularly brick-like, blocky…***Kadua*** p.p

43. Seeds strongly dorsiventrally compressed and winged; wing incurved or concave, margin entire or eroded; taxa exclusive to the Americas, mainly the Neotropics…44

43’. Seeds not strongly dorsiventrally compressed nor winged; or if slightly winged, exclusive to Polynesia, Asia, Africa or Australia 46

44. Herbaceous vine (except: *Manettia irwinii* Steyerm. with erect stems); axillary inflorescences, rarely terminal; capsule with septicidal dehiscence ***Manettia***

44’. Herbs, subshrubs, or shrubs, erect or decumbent; terminal inflorescences; capsule with loculicidal dehiscence, or loculicidal followed by septicidal dehiscence 45

45. Corolla hypocrateriform, red, orange, yellow, lilac, white, 7–45 mm long, internally glabrous, sparsely hairy in the lower half of the corolla tube, or ring of trichomes near the base of the tube; distributed from Mexico to Central America ***Bouvardia***

45’. Corolla infundibuliform, always white, 3–10 mm long, corolla lobes densely pubescent, corolla tube glabrous or sometimes with a fringe of trichomes near the base; endemic to Mexico…***Martensianthus***

46. Trigonous, or angular seeds…47

46’. Elliptic, ovoid, subglobose, navicular, cup-shaped, dorsiventrally compressed, cerebriform or crateriform seeds (with a broad ventral depression or concavity). 59

47. Mainly (sub)shrubs or small trees ca. 6 m tall 48

47’. Mainly herbs, sometimes with short woody underground stems…51

48. Fruits with septicidal dehiscence, usually followed by partial apical loculicidal dehiscence, resulting in two semi-split valves (diplophragmous capsules); seeds sometimes can be dorsiventrally compressed. ***Hedyotis***

48’. Fruits with only loculicidal dehiscence, or a combination of loculicidal and septicidal dehiscence; seeds irregular-angular or laterally compressed, but never dorsiventrally compressed…49

49. Corolla infundibuliform; fruit opened by loculicidal and septicidal dehiscence into 4 diverging valves…***Agathisanthemum***

49’. Corolla hypocrateriform; fruit opened by loculicidal dehiscence, or tardily septicidal, never separating into 4 diverging valves…50

50. Corolla with appendaged lobes…***Kadua*** p.p.

50’. Corolla without appendaged lobes…***Conostomium***

51. Capsular fruit which splits loculicidally and septicidally into 4 diverging valves…***Dibrachionostylus***

51’. Capsular fruit which splits only loculicidally from apex…52

52. Monomorphic flowers (only short-styled flowers); anther and stigma included in corolla tube; corolla always long tubular…53

52’. Dimorphic flowers (long-styled flowers and short-styled flowers); variety in corolla shape 54

53. Bifid stigma; seed exotesta alveolate to reticulate-alveolate with tangential walls never punctate; pollen 3-apertured, sometimes 4–5 (rarely 6); mesopore surrounded by a ring; short colpus often with fish tail endings as endoaperture; mostly with double reticulum….***Kohautia***

53’. Stigma capitate-bilobed; seed exotesta reticulate with prominent radial walls, tangential walls slightly to densely punctate; pollen 4–6-apertured (except *Cordylostigma virgatum* (Willd.) Groeninckx & Dessein, 3-apertured); mesopore without ring; endocingulum as endoaperture; reticulum simple…***Cordylostigma***

54. Leaves ovate to oblong and with somewhat decurrent leaf base, the uppermost appearing whorled; corolla lobes spreading to reflexed in fresh flowers, giving corolla a star-shaped appearance in some species…***Debia***

54’. Leaves variable in shape; corolla lobes not reflexed, and not appearing star-shaped…55

55. Flowers always heterostylous arranged in terminal inflorescences; pollen grains with endocolpus or endocingulum, and a mesoporus surrounded by a costa at the inside of the grain (compound ora)…***Lelya***

55’. Flowers heterostylous or homostylous, arranged in inflorescences terminally or axillary (or pseudoaxillary); pollen grains with endocingulum or lalongate endocolpus (simple ora). 56

56. Seeds irregularly angular with 3–5 pits/depressions on either side of the seed…***Involucrella***

56’. Seeds trigonous without pits/depressions on its surface 57

57. Corolla tube always glabrous; subglobe stigma; pollen with endocingulum as endoaperture ***Edrastima***

57’. Corolla tube pubescent or glabrous; bifid stigma; pollen with lalongate endocolpus as endoaperture 58

58. Flowers always homostylous; stamens and style always exserted…***Scleromitrion***

58’. Flowers homostylous or heterostylous; stamens and style usually included…***Oldenlandia***

59. Small herbs up to 10-30 cm tall, mainly prostate, creeping, weakly ascending, or erect. 60

59’. Subshrubs, shrubs or small trees, if herbs, more than 30 cm tall; mainly erect, decumbent, or scandent. 66

60. Axillary inflorescences; exclusively from the Americas or Australia 61

60’. Terminal inflorescences; exclusively from Africa and Madagascar. 64

61. Leaves narrowly oblanceolate or very narrowly elliptic; corolla rotate, marcescent on fruit; corolla tube short; stamens with filaments attached to base of corolla as well as to ovary; exclusively from Australia ***Synaptantha***

61’. Leaves broadly ovate, elliptic, suborbicular, or subrhombic; corolla infundibuliform or subhypocrateriform, not marcescent on fruit; corolla tube long; stamens with filaments attached only to base of corolla; exclusively from the Americas. 62

62. Plant spreading, erect to prostate, with tubers ca. 7 mm wide; calyx 6-8-lobate; seeds boat-shaped, cymbiform ***Lucya***

62’. Plant creeping without tubers; calyx 4-lobate; seeds oblate, ellipsoid, rhomboidal, depressed-subgloboid, obtusely angulate, or strongly compressed. 63

63. Flowers homostylous; corolla subhypocrateriform; capsule narrowly turbinate or narrowly obconic, dehiscing loculicidally and later separating into four narrow segments…***Oldenlandiopsis***

63’. Flowers heterostylous; corolla infundibuliform; capsule obovate, dehiscing loculicidally without separation of different segments…***Terrellianthus***

64. Ovary semi-inferior to superior (secondarily semi-inferior); capsule widely triangular, dehiscing loculicidally followed by septicidal dehiscence in the apex, into four diverging valves; beak (ovary roof) very distinct…***Mitrasacmopsis***

64’. Ovary notoriously inferior; capsule elliptic, obovoid, ovoid, rarely subglobose, dehiscing loculicidally, or loculicidal and septicidal, but never separating into four valves; beak indistinct…65

65. Plants succulent, mainly prostrate; flowers with stamens and style both exserted; calyx always 4-lobate; ovary with many ovules per locule; seeds with ventral groove absent; pollen 3-colporate…***Phialiphora***

65’. Plants not succulent, mainly erect; flowers with stamens and style both included; calyx 2-4-lobate;ovary with few ovules per locule (less than 10); seeds with ventral groove present; pollen 5-6 colporate…***Astiella***

66. Herbs or shrubs, lianescent, climbing and scandent…67

66’. Herbs, subshrubs, shrubs, or small trees, erect, prostrate, creeping, decumbent…68

67. Capsule dehiscing loculicidally and septicidally into 4 diverging valves; seeds discoid, convex dorsally, flattened ventrally; pollen 3-colporate; distributed in Africa (Tanzania) ***Pseudonesohedyotis***

67’. Capsule dehiscing loculicidally from apex followed by partial septicidal dehiscence, but not separating into valves; seeds dorsiventrally compressed, convex or saucershaped, often with narrowly winged margin, with a raised central hilar ridge; pollen 4-(5)-colporate; distributed in Tropical Asia…***Dimetia***

68. Seeds mainly with a hilar ridge in a ventral deep or shallow depression (crateriform, cymbiform to shallowly cupshaped seeds)…69

68’. Seeds without a hilar ridge in a ventral depression…70

69. Pollen grains (5-)6-12-aperturate; exclusively from Asia ***Neanotis***

69’. Pollen grains 3-(4)-aperture; exclusively from the Americas…***Houstonia***

70. Shrubs to small trees (ca. 6 m tall); corolla with appendaged lobes. ***Kadua***

70’. Herbs more than 30 cm tall, to (sub)shrubs; corolla without appendaged lobes…71

71. Leaves suborbicular, ovate, oblong to oblanceolate, widely elliptic, or narrowly lanceolate 72

71’. Leaves filiform, linear, narrowly elliptic 74

72. Plants usually glabrous throughout; capsule usually winged; seeds appearing pitted due to shallow depressions bordered by thick and sinuate walls; distributed to tropical and subtropical Asia to Pacific ***Leptopetalum***

72’. Plants glabrescent, minutely puberulent or pubescent; capsule not winged; seeds without pitted appearance; distributed to the Americas. 73

73. Plants growing in high-altitude grasslands (1500–3000 m); capsule with septicidal dehiscence, sometimes only the beginning of the dehiscence loculicidal. ***Arcytophyllum***

73’. Plants growing in moist soil, shaded places, banks, slopes, among rocks, sides and bases of cliffs, montane rain forest (330-1430 m); capsule with first loculicidal then septicidal dehiscence ***Mexotis***

74. Axillary inflorescences, in a spiciform arrange ***Manostachya***

74’. Terminal inflorescences, variable…75

75. Homostylous flowers, some heterostylous. 76

75’. Always heterostylous flowers…78

76. Stamens and style exserted in homostylous flowers; heterostylous flowers can be present; corolla tube short; capsule globose; exclusively from Africa ***Amphiasma***

76’. Stamens and style included; always homostylous flowers; corolla tube well developed; capsule subglobose, depressed ovoid, generally longer than broad, laterally compressed; exclusively from Australia 77

77. Corolla lobes geniculate with a distinct transverse line of hairs at knee or corolla lobes spathulate; seeds generally cerebriform, broadly sulcate, rarely scutelliform, with hilum near centre of ventral surface, rarely on conspicuous central ridge ***Dolichocarpa***

77’. Corolla lobes not geniculate; without a distinct transverse line of hairs part way along corolla lobes, with a line of hairs at base of corolla lobes, corolla lobes not spathulate; seeds scutelliform, broadly elliptic or oblong in outline, with hilum centrally placed on a prominent ventral ridge ***Paranotis***

78. Capsules dehiscing loculicidal and septicidal into 4 diverging valves; locules with reduced number of seeds (5–10). ***Hedythyrsus***

78’. Capsules dehiscing only loculicidal, sometimes followed by partial septicidal dehiscence, but never separating into four valves; locules with numerous number of seed (up to 47 per capsule). 79

79. Hypocrateriform corolla; corolla tube 12-37 mm long. ***Pentanopsis***

79’. Infundibuliform or hypocrateriform corolla; corolla tube 1-10 mm long…80

80. Stamens inserted at two levels in the corolla tube ***Amphistemon***

80’. Stamens inserted at the same levels in the corolla tube…81

81. Much-branched shrub up to 1.5 m tall; capsules broadly obovate to transversely broadly obovate; seeds winged; endemic from Madagascar ***Thamnoldenlandia***

81’. Perennial herbs or low shrubs, up to 50 cm tall, with or without woody tap root; capsules slightly compressed, turbinate, obovoid, ellipsoid, or subglobose; seeds not winged; exclusively from the Americas…82

82. Seeds ventral face with a punctiform centric hilum on flat, laterally compressed, or convex to rounded with hilar ridge; chromosome number x=13; endemic from Baja California, Mexico…***Stenotis***

82’. Seeds ventral face with a punctiform centric hilum on flat, or slightly convex surface, without hilar ridge; chromosome number x=9 or 10; distributed in southwestern USA or Mexico…***Stenaria***

## 5. Conclusion

### Toward a stable classification of Spermacoceae

Our plastome phylogeny represents a substantial advance over previous molecular studies of Spermacoceae, including for the first time chloroplast genome data for 127 accessions, representing 54 genera (64 %). Compared with earlier Sanger-based phylogenies, plastome-scale sequencing increased the number of aligned nucleotide sites available for phylogenetic inference by approximately 40-fold, resolving most major relationships within the tribe with strong support and providing the most comprehensive phylogenetic framework for the tribe to date. Genera such as *Lucya*, *Martensianthus*, and *Mexotis* were included for the first time in a phylogenetic analysis. Also, the phylogenetic positions of *Oldenlandiopsis* and *Terrellianthus* were clarified. Nevertheless, some backbone relationships remain weakly supported, particularly where successive divergences are separated by very short internal branches. Because the plastome is inherited as a single, non-recombining locus, it provides only a single genealogical history and cannot, by itself, distinguish among the evolutionary processes responsible for such patterns. A comparison with a nuclear ribosomal ITS phylogeny showed a topology that was largely congruent with the plastome tree, with most major clades recovered in both datasets, although several groups were resolved in conflicting positions between the two trees. Likewise, within rapidly diverging lineages such as *Hedyotis* and *Kadua*, complete plastome sequences appear to provide insufficient phylogenetic signal to fully resolve relationships among closely related species. Phylogenomic data from hundreds of low-copy nuclear loci, such as those generated using the Angiosperms353 target-capture system (Johnson et al. 2019), should improve resolution of these short backbone branches while allowing hybridization, introgression, and other sources of gene-tree discordance to be evaluated explicitly. Combined with broader taxon sampling, particularly in unresolved complexes such as Houstonia–Stenaria and Galianthe–Carajasia–Schwendenera, and *Spermacoce*, such data will further refine the classification of Spermacoceae and deepen our understanding of the evolutionary history of one of the largest and most taxonomically challenging tribes in Rubiaceae.

## CRediT authorship contribution statement

**Mariela Nuñez Florentin:** Conceptualization, Investigation, Formal analysis, Data curation, Visualization, Writing. **Kieran Claypool, Nusrat Huda, Katherine Green,** and **Greg Monzel:** Investigation (DNA extraction and library preparation). **Peter Schafran:** Methodology, Supervision. **Suman Neupane:** Conceptualization, Methodology, Formal analysis, Data curation, Funding acquisition, Project administration, Writing.

## Acknowledgements

We thank herbarium curators and staff at the Instituto de Botánica del Nordeste (CTES), Harvard University Herbarium (HUH), Missouri Botanical Garden (MO), New York Botanical Garden (NY), Jardim Botânico do Rio de Janeiro (RB), Smithsonian Institution (US) for deposition of, and access, to herbarium specimens used in this work. We also thank Javier E. Florentin and Roberto M. Salas.

